# Strawberry phenotypic plasticity in flowering time is driven by interaction between genetic loci and temperature

**DOI:** 10.1101/2023.11.29.569202

**Authors:** Alexandre Prohaska, Aurélie Petit, Silke Lesemann, Pol Rey-Serra, Luca Mazzoni, Agnieszka Masny, José F. Sánchez-Sevilla, Aline Potier, Amèlia Gaston, Krzysztof Klamkowski, Christophe Rothan, Bruno Mezzetti, Iraida Amaya, Klaus Olbricht, Béatrice Denoyes

**Affiliations:** Univ. Bordeaux, INRAE, Biologie du Fruit et Pathologie, UMR 1332, F-33140, France; INVENIO, MIN de Brienne, 110 quai de Paludate, 33800 Bordeaux, France; Hansabred GmbH & Co. KG, Dresden, Germany; Department of Agricultural, Food and Environmental Sciences, Marche Polytechnic University, 60131 Ancona, Italy; National Institute of Horticultural Research, Konstytucji 3 Maja 1/3, 96-100 Skierniewice; Centro IFAPA de Málaga, Instituto Andaluz de Investigación y Formación Agraria y Pesquera (IFAPA), 29140, Málaga, Spain; Unidad Asociada de I+D+i IFAPA-CSIC Biotecnología y Mejora en Fresa, 29010, Málaga, Spain

**Keywords:** flowering time, genotype × environment interaction (G×E), phenotypic plasticity, QTL-by-Environment Interaction (QEI), Quantitative Trait Locus (QTL), strawberry

## Abstract

The flowering time, which determines when the fruits or seeds can be harvested, is known to be sensitive to plasticity, i.e. the ability of a genotype to display different phenotypes in response to environmental variations. In the context of climate change, strawberry breeding can take advantage of phenotypic plasticity to create high-performing varieties adapted either to local conditions or to a wide range of climates. To decipher how the environment affects the genetic architecture of flowering time in cultivated strawberry (*Fragaria ×ananassa*) and modify its QTL effects, we used a bi-parental segregating population grown for two years at widely divergent latitudes (5 European countries) and combined climatic variables with genomic data (Affymetrix® SNP array). We detected 10 unique flowering time QTL and demonstrated that temperature modulates the effect of plasticity-related QTL. We propose candidate genes for the three main plasticity QTL, including *FaTFL1* which is the most relevant candidate in the interval of the major temperature-sensitive QTL (6D_M). We further designed and validated a genetic marker for the 6D_M QTL which offers great potential for breeding programs, for example for selecting of early-flowering strawberry varieties well adapted to different environmental conditions.

**Highlights:** A GXE study of a segregating strawberry population in Europe showed that temperature is the main driver of flowering time plasticity. A genetic marker was designed for the main QTL.

## Introduction

Phenotypic plasticity describes the ability of a given genotype to produce distinct phenotypes in response to different environments (Pigliucci, 2005). It allows species, populations, or genotypes to cope with rapid environmental changes, including global climate change. In crop species, knowledge of trait plasticity is an important element of the success of a variety. The breeder can select locally adapted varieties which, by taking advantage of local conditions, will give better results than widely adapted varieties (Ceccarelli, 1989; Kusmec et al., 2018).

The central approach to characterize plasticity of a trait is to identify for each genotype the reaction norm, which describes how the target phenotype of a specific genotype varies as a function of the environmental variables to which the genotype is exposed (Sultan, 1987). Genotype-by-environment interactions (G×E) are observed when reaction norms are non-parallel between genotypes (Pigliucci, 2005). To assess this interaction, multiple genotypes or populations must be studied in a large range of environments. Numerous statistical approaches have been developed to study this interaction (reviewed in Li et al., 2017). G×E can be detected by an ANOVA with fixed or mixed models but the interpretation of the interaction is limited with these approaches. Other approaches such as AMMI or joint regression model allow the estimation of plasticity parameters to explain the interaction. Factorial regression allows the inclusion of explicit environmental factors (i.e. covariates) in G×E models along with a direct evaluation (i.e. quantification) of the importance of these covariates for G×E explanation (Malosetti et al., 2013; Lombardi et al., 2022). As a consequence, this model makes it possible to identify the environmental parameters that are biologically relevant to the trait.

One of the traits described as highly sensitive to plasticity is flowering time (Blackman, 2017). It is a trait critical for the adaptation of a variety to a particular region, as it determines when fruits or seeds are harvested and the final yield. It is regulated by endogenous genetic components and environmental factors (Cho et al., 2017). Strawberry (*Fragaria ×ananassa*), the most cultivated berry worldwide with a total harvested area of 389,665 ha and a total production of 9,175,384 T in 2021 (FAOSTAT, https://www.fao.org/faostat/en/#data), is widely grown in the northern and southern hemispheres. New varieties adapted to a wide range of latitudes, from tropical/subtropical to cold temperate climates, have been selected using different strategies (Senger et al., 2022). Varieties cover more or less restricted regions: for example, ’Fortuna’ is grown in Florida (USA) but also in Mexico, Spain, Egypt and Morocco and, conversely, ’Florence’ is restricted to Norway. As most breeding programs are organized according to seasonality, the genetic architecture of flowering time and its plastic responses to environments need to be characterized (Li et al., 2018). Unlike perpetual flowering (PF) mutants where flowers are initiated continuously, flowering in seasonal flowering (SF) genotypes occurs in spring and is the consequence of floral initiation that occurred the previous autumn under low temperature and short days (Gaston et al., 2020, 2021). Thus, after dormancy, plant growth resumes in spring and inflorescences initiated the previous year emerge and flower. While the genetic and molecular control of floral initiation has been extensively studied in diploid species (Iwata et al., 2012; Koskela et al., 2012; Gaston et al., 2020 and 2021) and, more recently, in cultivated octoploid strawberry (Nakano et al., 2015; Koembuoy et al., 2020; Gaston et al., 2021; Muñoz-Avila et al., 2022), studies on the genetic control of flowering time are scarce and focus exclusively on diploid species (Samad et al., 2017).

Exploring how flowering time and its phenotypic plasticity are genetically and environmentally controlled is essential for breeding better adapted strawberry varieties. To achieve this objective, we built a concerted European project (GoodBerry) to study the response of flowering time to diverse environments in a bi-parental segregating population cultivated at very different latitudes (5 southern and northern European countries) over a two-year period. The integration of strawberry genomic data (Affymetrix® SNP array) with phenotypic and climatic data enabled us to detect three flowering time quantitative trait loci (QTL) for which the overall mean flowering times co-localized with plasticity parameters. We further designed and validated a genetic marker for the main highly temperature-sensitive QTL (6D_M), which offers strong potential for selecting strawberry varieties well adapted to different climates.

## Material and methods

### Plant material and phenotyping

A pseudo full-sibling F1 population of 109 individuals derived from two SF varieties with contrasting European cultivation areas was obtained: ‘Candonga’ is widely cultivated in south of Spain and ‘Senga Sengana’, originally selected in Germany, is commonly grown in Poland. Nine experiments were conducted in five countries from north and south Europe in 2018 (5 experiments) and 2019 (4 experiments): Skierniewice, Poland (PL) (51°95’N); Dresden, Germany (GE) (51°05’N); Agugliano, Italy (IT) (43°32N); Douville, France (FR) (45°59’N); and Huelva, Spain (SP) (37°24’N) (Fig. 1A). To homogenise the physiological development of daughter plants of all individuals and parents, young plants obtained from a single nursery were sent to the five locations for plantation in 2017 and 2018 (Supplementary Fig. S1). For each individual of the genetic segregating population, 10 plants were grown in open field or under plastic tunnels, except in France where they were grown in soil-free pine bark substrate under plastic tunnel. Flowering time was defined as the date of observation of the first flower at anthesis.

**Figure 1.**
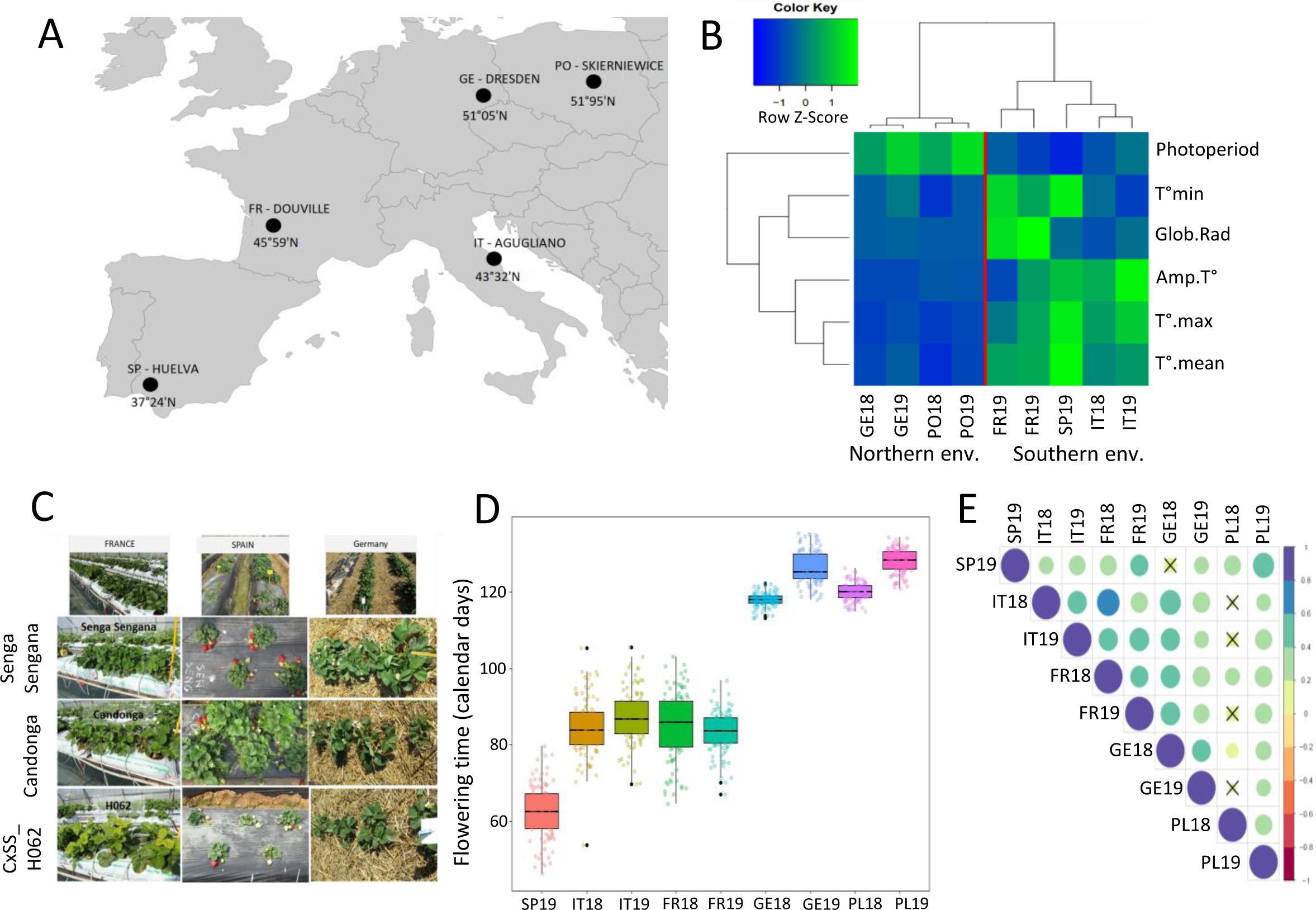
Environment description and flowering time of the ‘Candonga’ x ‘Senga Sengana’ segregating population in nine environments. (A) Location and latitudes of the five countries in which experimental trials were carried out. (B) Clustering of the nine experimental environments according to the six environmental covariates measured from the 1^st^ of January until end of flowering. (C) Three genotypes, ‘Candonga’, ‘Senga Sengana’ and ‘H062’ under three environments in 2018 (France, Spain and Germany). (D) Box plots showing the flowering time (calendar days) for the nine environments. (E) Pairwise Spearman correlation values (r) between flowering time (calendar days) in the nine environments. The r values are represented by coloured circles whose size varies according to their value. Crossed-out circles indicate non-significant correlations (*p*-value > 0.05). SP, Spain, IT, Italy, FR, France, GE, Germany, PL, Poland. 2018 and 2019 (18 and 19 respectively), years of experimentations. Sites are ordered by increasing order of latitude.

At each location, we recorded daily climatic variables, temperatures (mean, maximum, minimum; in °C) and global radiations (in kw/m²). These data were analysed by hierarchical clustering using Euclidean distance and Ward’s method. The distance was calculated based on environmental parameters: temperatures (mean, maximum, minimum, difference minimum-maximum per day), photoperiod, global radiation and sum of Growing Degree-Days (GDD) of the nine environments.

### Modeling flowering time

To model flowering time phenology, we tested four thermal time models: Growing Degree-Days (GDD, Wang, 1960), triangular (Hänninen, 1990a), sigmoid (Hänninen, 1990b) and Wang (Wang and Engel, 1998). These models assume that there is a relationship between the phenological stage (the flowering period) and the cumulative temperature above a threshold (base temperature, Tb or Tmin) over a given period. Temperature is expressed here in degrees Celsius (Chuine et al., 1998). This sum (SStar) is calculated from the starting date, t0. In addition, triangular and Wang models consider optimal (Topt) and maximal (Tmax) temperatures. In addition, to study the efficiency of predicting flowering date as a function of photoperiod or global radiation, we adapted the calculation of the GDD and triangular models to these two climatic parameters. Process-based models were adjusted by minimizing the residual sum of squares with the simulated annealing algorithm of Metropolis (Chuine et al., 1998) using the Phenology Modeling Platform software (PMP5; http://www.cefe.cnrs.fr/fr/recherche/ef/forecast/phenology-modelling-platform) (Chuine et al., 2013). Adjustment was repeated 20 times to ensure that the global optimum had been reached. To simulate the flowering time, we included data from both parents and from 102 individuals for whom all the data for the nine environmental conditions were available. For further analyses, we retained the most parsimonious model and the best efficiency (R^2^).

### Statistical modeling for variance components and heritability estimation

To study the variation in flowering time (GDD) in our segregating population, we fitted a linear mixed-effects model (LMM) by maximum-likelihood (LME4 package; Bates et al., 2015) following equation 1 (eqn1):

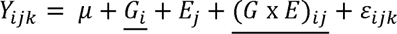

where µ is the overall mean of the population, Gi the random effect of genotype i, Ej the fixed effect of environment i, G x Eij the random interaction effect between genotype i and environment j and eijk the residual term assumed to be normally distributed. The best sub-model was selected according to Fisher test for the environment effect log-likelihood ratio tests (LRT) for random effects with the lmerTest R package (Bates et al., 2021). The selected model was re-fitted by Restricted Maximum Likelihood with the plantTrialLmmFitCompSel function from the rutilstimflutre R package (Timothee Flutre’s personal R code. URL https://github.com/timflutre/rutilstimflutre).

The broad-sense heritability at the whole design level (H2) was derived from the variance components of eqn1 and calculated in equation 2 (eqn2):

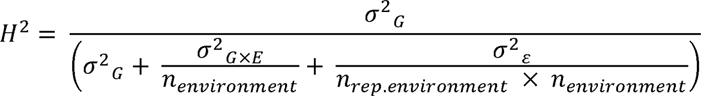

with genotype (G) variance (σ^2^) at the numerator. Random variance components involving environment (E) were divided by the number of environments (n _environments_). Residual variance was divided by the number of environments multiplied by the average number of replicates per environment (n _rep.environment_).

### Statistical modeling of plasticity parameters

We performed complementary statistical approaches to compute genotype specific plasticity parameters using the additive main effects and multiplicative interaction (AMMI) method, the joint regression analysis also named Finlay-Wilkinson (FW) regression model and the factorial regression model.

i. The additive main effects and multiplicative interaction (AMMI) method combines analysis of variance (ANOVA) to model main effects of genotype and environment and principal component analysis (PCA) to decompose the complex structure of G×E into Interactive Principal Component Axes (IPCA) (Gauch, 2013) (equation 3, eqn3):

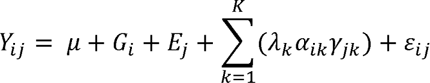

where Y_ij_ is the mean phenotypic performance of genotype i in environment j; µ is the intercept; G_i_ the fixed effect of genotype i; e_j_ is the fixed effect of environment j; λ_k_ is the singular value for the IPCA k; α_ik_ are the genotypic IPCA scores and γ_*jk*_ the environmental loadings for axis k; ε_ij_ is the residual term G×E not captured by the model and some error deviation.

We derived AMMI Stability Value (ASV) from eqn3 for each genotype as the relative influence of IPCA1 and IPCA2 scores based on their interaction sum of squares (SS) according to Purchase (1997) using the formula:

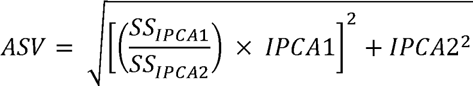

where (SS_IPCA1_/SS_IPCA2_) is the weight assigned to the IPCA1 value by dividing the IPCA1 SS by the IPCA2 SS; IPCA1 and IPCA2 scores were the genotypic scores derived from the AMMI model. A large positive ASV value indicates a genotype that is adapted to particular environments. A small (close to zero) ASV value indicates a stable genotype across environments (Bakare et al., 2022).

(ii) In the joint regression analysis (FW regression), G×E is modeled by regressing mean phenotypic performance of genotypes on an environmental index. The index value of each environment is calculated as the mean of all individuals of the flowering time in that environment (Finlay and Wilkinson, 1963). Then, the intercept and slope for each genotype are calculated by regressing genotypic performance on the environmental index as in equation 4 (eqn4):

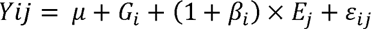

where µ+G_i_ represents the average performance of a genotype considering all environments; the slope 1+ β_i_ represents the regression coefficient of the model and is the linear response to environment; the residual variance of the term ε, which measures the scatter of points about the regression lines, represents the non-linear response to environment (non-linear plasticity).

(iii) The factorial regression model allowed the description of G×E by using explicit covariates as environmental factors (Tmean, Tmin, Tmax, GDD, photoperiod or global radiation). Each climatic covariate was tested successively at a significance threshold of 5% to be incorporated into the following equation 5 (eqn5):

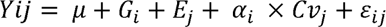

where the genotypic response of genotype i in environment j is described through its sensitivity *α_i_* to the tested covariate *Cv_j_*. Slopes from eqn4 and eqn5 were computed with the script adapted from Diouf et al. (2020).

### Other statistical analyses

Correlations between the mean flowering times were performed with “rcorr” procedure of the Hmisc R package (https://cran.r-project.org/web/packages/Hmisc/Hmisc.pdf) and a Bonferroni correction was applied at a threshold of 5%. Pairwise comparisons were performed using Student’s T-test (*p* < 0.05).

### Development of linkage maps

Single dose markers (SD) from the Affymetrix® array (Hardigan et al., 2020) that were in backcross configuration and segregated 1:1 (Rousseau-Gueutin et al., 2008) were used for genetic map construction using JoinMap® 5.1 software (Van Ooijen, 2011). Grouping was performed using independence log of the odds (LOD) and the default settings in JoinMap®. Linkage groups (LG) were chosen from a LOD higher than 10 for all of them. Map construction was performed using the maximum likelihood (ML) mapping algorithm and the parameters described in Labadie et al. (2022). Mapping results are displayed using MapChart (Voorrips, 2002).

### QTL mapping and QTL-by-environment interactions (QEI)

The female and male linkage parental maps based on the 109 individuals were used separately for QTL analysis. Flowering time expressed as GDD by environment and plasticity parameters (i.e. ASV and IPCA values from AMMI model, slopes and residual variances from joint and factorial regressions), represented the phenotypic data. QTL detection was performed by simple interval mapping (SIM) using R/QTL package (Broman et al., 2003). Permutation analysis (1000 permutations) was performed to calculate the critical LOD score. QTL with LOD values higher than the LOD threshold at *p* ≤ 0.05 were considered significant. When one QTL was found significant, we used composite interval mapping (CIM) with one co-variable at the position of the significant QTL and reiterated the analysis until no new significant QTL were detected. Bayesian credible interval was calculated using the function ‘bayesint’ at probability of 0.95. The proportion of phenotypic variance explained by a single QTL was calculated as the square of the partial correlation coefficient (R^2^).

We searched for QTL × temperature by SIM following a two-step procedure by testing the temperature as an interactive (eqn6)

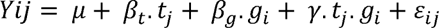

and then as an additive (eqn7) covariate:

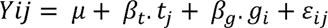

where Yij is the trait value for allele *i* (*i* = 1,2) in environment *j* among the nine location-by-year combinations (overall mean of the flowering time); *t_j_*, the mean temperature in environment j; *g_i_*, the QTL effect for genotype i; γ, the QTL × temperature interaction coefficient; *ε_ij_*, the residual term. Evidence of QEI was assessed by taking the LOD difference between equations 6 (eqn6) and 7 (eqn7).

### Candidate genes

Candidate genes likely to play a role in flowering time were identified into the Bayesian credible intervals common to the different QTL detected in each region showing the highest number of significant QTL: 3A_M, 6A_M and 6D_M. Homologs of known flowering time genes were selected as candidate genes in ‘Camarosa’ (Edger et al., 2019) and ‘Royal Royce’ (Hardigan et al., 2021; https://phytozome-next.jgi.doe.gov/info/FxananassaRoyalRoyce_v1_0) genomes.

### Marker design

We developed a subgenome-specific Kompetitive Allele Specific PCR (KASP) marker (Smith and Maughan, 2015) linked to the major 6D_M QTL. The Affymetrix® marker AX-184201950 localized in the middle of the QTL harbours a C/T SNP (Hardigan et al., 2020). Specific primer design was performed using BatchPrimer3 software (http://probes.pw.usda. gov/batchprimer). Genotyping was done on the segregating population and on additional 94 genotypes using the KASP procedures described by LGC Genomics (Supplementary Table S1). Genotyping data were viewed as a cluster plot (LightCycler® 480 qPCR software, Roche). The significance of the relationship between phenotype and genotype was determined using Wilcoxon test.

## Results

### Strawberry flowering time plasticity under natural conditions

We studied the flowering time of the ‘Candonga’ x ‘Senga Sengana’ strawberry bi-parental population cultivated in five countries covering a wide range of latitudes (Fig. 1A; Supplementary Table S2). Cultures were conducted under field (PL, GE, IT) or tunnel (FR, SP) environments. Flowering time was measured during two successive years 2018 and 2019 (hereafter named 18 and 19), except Spain, which was only measured in 2019, and thus nine location-by-year combinations. These nine environments clustered into two groups that overlapped southern (SP, IT, FR) and northern (GE, PL) areas in Europe (Fig. 1B).

The bi-parental population was issued from a cross between two varieties displaying geographical opposite cultural adaptation with ‘Candonga’ selected in Southern Europe and ‘Senga Sengana’ in Northern-Eastern Europe (Fig. 1C). Flowering time followed a latitude gradient when expressed as calendar days and showed a larger variation in southern environments than in northern ones (Fig. 1D). At Northern latitudes, the population flowered on average six to eight days earlier in 2018 than in 2019 (Fig. 1D). Notably, phenotypic correlations between environments were strictly positive but were weak (0.27-0.59) (Fig. 1E; Supplementary Table S3) suggesting genotype-by-environment interactions with changes in ranking.

### Growing Degree Days (GDD) for expressing the flowering time

In strawberry, temperature has been described as the main environmental factor affecting the flowering time (Le Mière et al., 1998; Opstad et al., 2011) whereas photoperiod has been reported to influence flowering time in PF genotypes (Sønsteby and Heide, 2007) or global radiation in SF genotypes (Krüger et al., 2022). We tested four models: GDD, triangular, sigmoid and Wang based on temperatures, global radiation and/or photoperiod. Whatever the model, the best efficiency was obtained with thermal times and Tmean (85%) (Table 1). Models were not improved by adding the effect of global radiation or photoperiod. Estimates of the parameters for each model were also similar for both the base temperature (Tb), -1.7—1.8 °C except for the Wang model (-13.6 °C), and the starting date (t0), January 1^st^. The triangular and Wang models gave in addition an optimum temperature (Topt) at 24-25 °C and a maximum temperature (Tmax) at 34-35 °C. This temperature was not reached under our conditions and we therefore retained the most parsimonious GDD model for further analysis.

**Table 1.** Models for predicting flowering date as a function of temperature, photoperiod and/or global radiation: GDD, Triangular, Sigmoid, Wang. For photoperiod and global radiation, calculation of the GDD and triangular models were adapted to these two climatic parameters. Efficiency, ratio (SStot-SSres/SStot); RMSE, root mean squared error; t0, starting date in calendar day; SStar, sum calculated by the model from t0; Nobs, number of observations; Tb, base temperature; Tmin, minimum temperature; Topt, optimum temperature; Tmax, maximum temperature; PPmin, PPopt, PPmax or GRmin, GRopt, GRmax for minimum, optimum, maximum of photoperiod or global radiation; Temperatures and global radiations are expressed in °C and Watt/m2 respectively. Models were calculated with the PMP5 software (http://www.cefe.cnrs.fr/fr/recherche/ef/forecast/phenology-modelling-platform). Parameters of the models were adapted to the range of values of the climatic factors, Tmean (Tm), Global radiation (GR) or Phtoperiod (PP). d and e are Sigmoid model parameters.

| Model | Efficiency (%) | RMSE | t0 | SStar | Nobs | Tb_Tmin / PPmin / RGmin | Topt / PPopt / RGopt | Tmax or PPmax / RGmax | d | e |
| --- | --- | --- | --- | --- | --- | --- | --- | --- | --- | --- |
| **Simple model** |  |  |  |  |  |  |  |  |  |  |
| *Temperature* |  |  |  |  |  |  |  |  |  |  |
| GDD:Tm | 0.85 | 9.07 | 1 | 824.7 | 918 | -1.7 |  |  |  |  |
| Triangular:Tm | 0.85 | 9.07 | 1 | 31.7 | 918 | -1.8 | 24.4 | 34.1 |  |  |
| Sigmoid:Tm | 0.85 | 9.00 | 1 | 31.4 | 918 |  |  |  | -0.19 | 11.32 |
| Wang:Tm | 0.85 | 9.00 | 1 | 30.4 | 918 | -13.6 | 25.0 | 34.3 |  |  |
| *Photoperiod* |  |  |  |  |  |  |  |  |  |  |
| GDD:PP | 0.37 | 18.63 | 1 | 237.2 | 918 | 8.0 |  |  |  |  |
| Triangular:PP | 0.62 | 14.47 | 1 | 47.50 | 918 | 7.8 | 12.10 | 14.09 |  |  |
| *Global Radiation* |  |  |  |  |  |  |  |  |  |  |
| GDD:RG | 0.07 | 22.59 | 66 | 12482 | 918 | 749.7 |  |  |  |  |
| Triangular:RG | 0.68 | 13.06 | 1 | 50.8 | 918 | 141.9 | 888.9 | 2681.2 |  |  |
| **Multiplicative models** |  |  |  |  |  |  |  |  |  |  |
| GDD:Tm and Triangular:PP | 0.85 | 9.05 | 1 | 528.5 | 918 | 0.2 | 13.2 | 24 |  |  |
| GDD:Tm and Triangular:RG | 0.79 | 11.00 | 16 | 198.7 | 918 | 0.5 | 1615 | 2470 |  |  |

We hypothesized that genotypes differed in the heat units necessary to trigger flowering. Therefore, we calculated the GDD value of each individual with parameters t0 as the 1^st^ of January and Tb as -1.7°C. We further plotted reaction norms for flowering time, expressed as calendar days (Fig. 2A) or GDD (Fig. 2B), for all individuals and parents across the environmental gradient quantified by the population means of calendar days or GDD.

**Figure 2.**
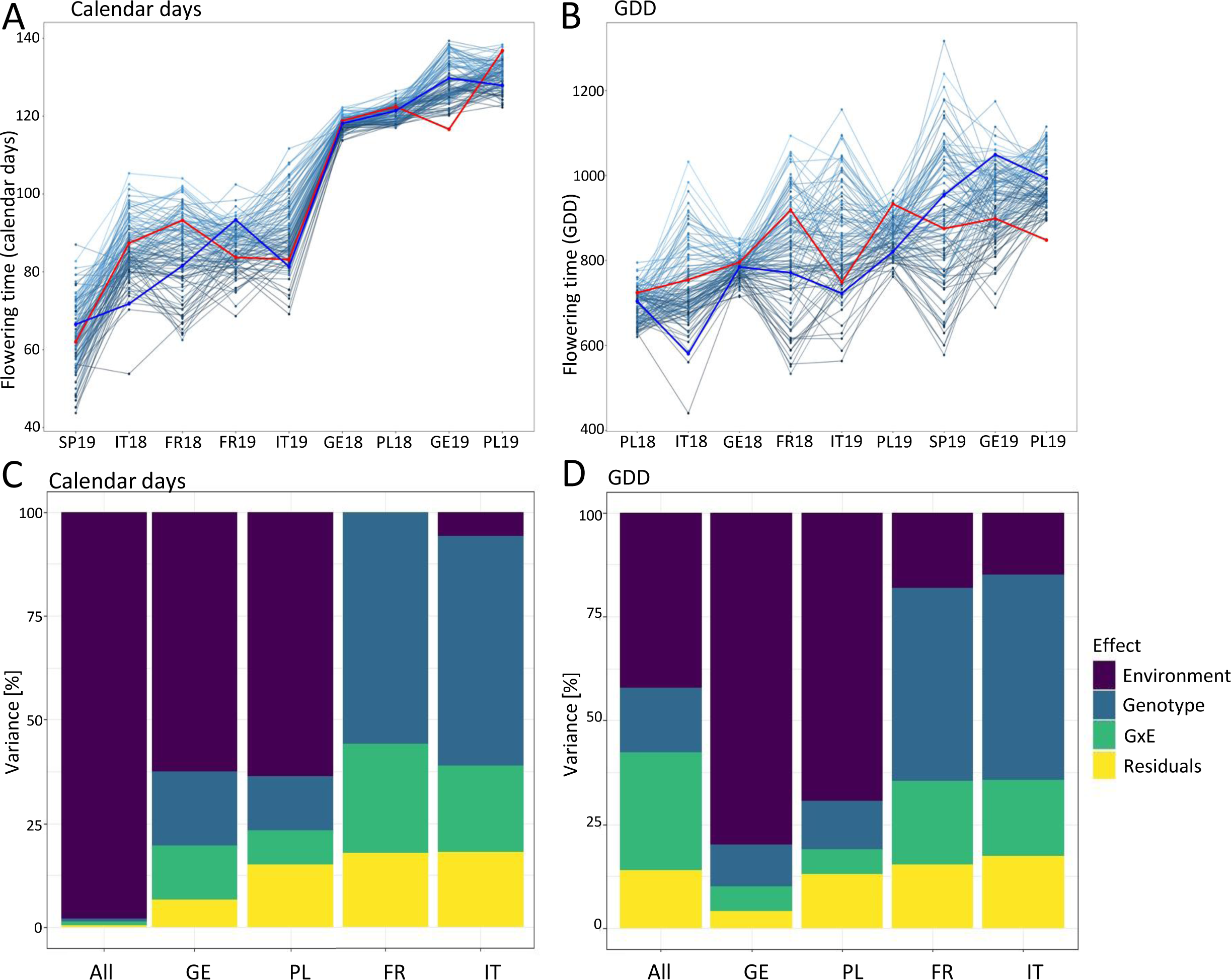
Reaction norms and analyses of variance for flowering time. (A-B) Reaction norm for flowering time expressed in calendar days (A) or Growing Degree-Days (GDD) (B). The nine environments are ranked in increasing order of flowering time. Each line connects the flowering time values of individuals across environments. Red and blue lines represent ‘Candonga’ and ‘Senga Sengana’ respectively. (C-D) Variance partitioning of flowering time (calendar days (C) and GDD (D)) by the linear mixed model for the nine environments or for each country. SP, Spain, IT, Italy, FR, France, GE, Germany, PL, Poland. 2018 and 2019 (18 and 19 respectively), years of experimentations.

The general linear mixed-effects model (eqn1) revealed that at the whole design level and at the country level, all effects (Genotype, Environment, G×E) were significant (Supplementary Table S4). At this whole design level, the environment, the factor contributing most to phenotypic variance, was more important when flowering time was expressed as calendar days (Fig. 2C) rather than as GDD (Fig. 2D). The proportions of G×E and Genotype variances of flowering time increased substantially towards Southern environments (SP, for which a single year of study could be performed, was not included in this analysis). The G×E variance was further split into a most contributing Genotype by Location (GxL) term and a significant Genotype by Year interaction (G×Y) term (Supplementary Table S4).

By-site heritabilities for both calendar day and GDD were higher in Spain (H² = 0.94, 0.95), Italy (H² = 0.92, 0.89) and France (H² = 0.91, 0.88) than in Northern countries, Germany (H² = 0.59, 0.32) and Poland (H² = 0.40, 0.34) (Supplementary Table S5). In subsequent analyses, we have retained the data relating to the flowering period expressed in GDD, as they summarise the data with high efficiency by clearly identifying the heat demand of the plants for flowering, while the calendar days reflect a combination of multi environmental factors.

### Plasticity parameters involved in G×E

For each model, AMMI, joint (FW) regression and factorial regression, analyses of variance revealed significant genotype, environment and G×E effects (*p* < 0.001) (Supplementary Tables S6, S7, S8). We further characterized G×E at the genotypic level with plasticity parameters derived from the three models (i.e., AMMI, FW and factorial regression) subsequently used for QTL mapping.

### AMMI

Decomposition of the genotype-by-environment interaction through the AMMI model (Supplementary Table S6) showed a large number of significant IPCA values (from IPCA1 to IPCA9) (*p* < 0.01) using the F test of Gollob (1968). Each of these nine IPCA values explained from 4.3% to 21% of the variation in the SS_GxE_, disclosing the complexity of the interaction patterns. The first two components captured less than half of the original variance (36.4%) with most of the environments poorly represented, complicating the interpretation of the biplots (Supplementary Fig. S2). The AMMI stability value (ASV) calculated on the first two IPCA ranged from 0.06 to 1.30 across the 109 individuals and the two parents (Fig. 3A, Supplementary Table S9). The genotypes ‘H091’, ‘H0104’ and ‘H077’ had the lowest ASV values, while the genotypes ‘H027’, ‘H073’ and ‘H122’ had the highest values.

**Figure 3.**
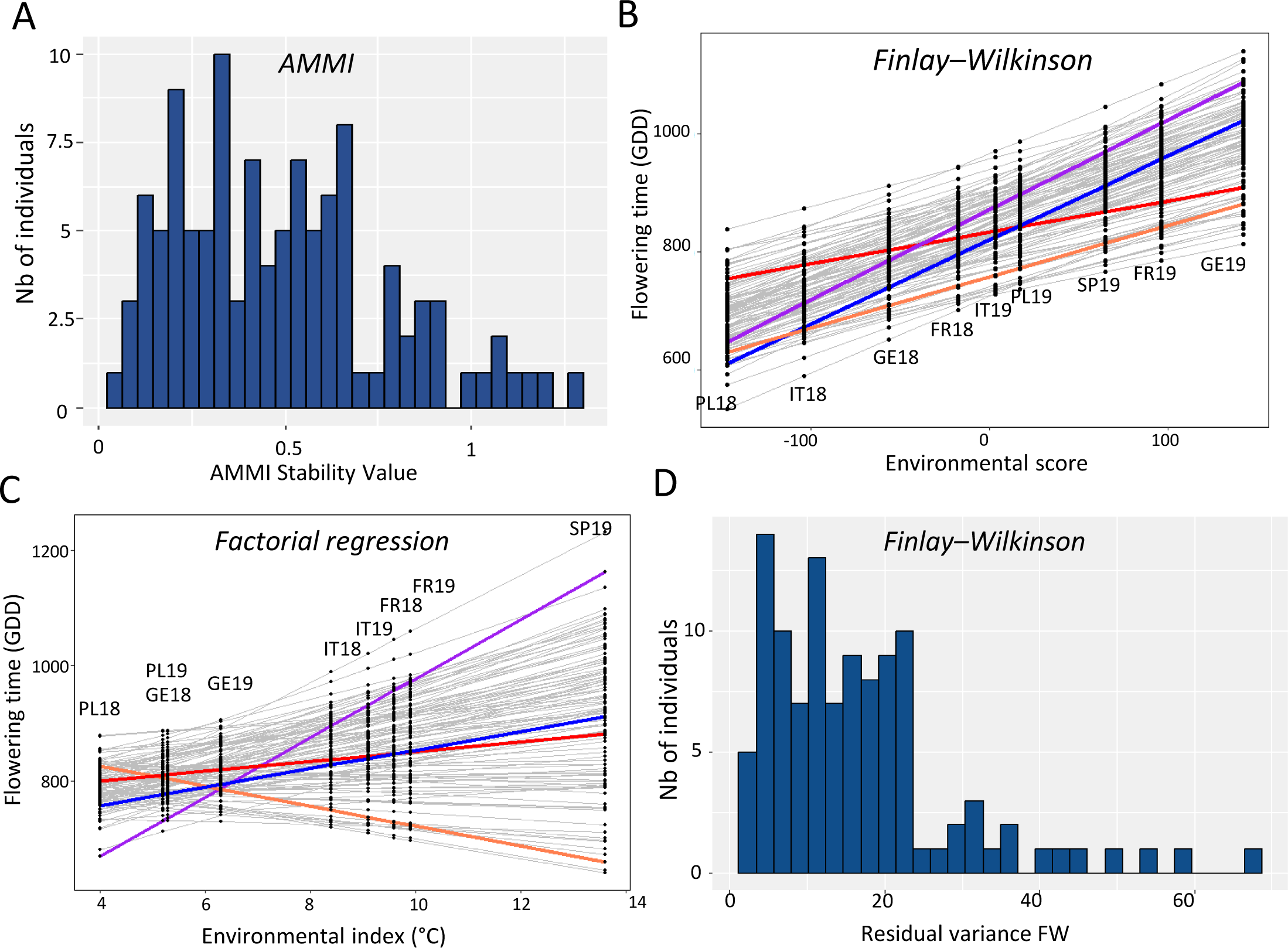
Distributions of plasticity parameters for all individuals and parents in the segregating population for flowering time. (A) Histogram distribution of AMMI Stability Values (ASV) calculated for each genotype as the relative influence of IPCA1 and IPCA2 scores based on the sum of squares of their interaction. (B-C) Slopes with Finlay–Wilkinson and factorial regression models for flowering time (GDD). Regressions of phenotypic performances of genotypes on environmental index (B) or on mean temperature (C). ‘Candonga’, ‘Senga Sengana’, ‘H102’ (example of a late flowering genotype) and ‘H056’ (example of an early flowering genotype) are plotted as red, blue, purple and orange lines, respectively. SP, Spain, IT, Italy, FR, France, GE, Germany, PL, Poland. 2018 and 2019 (18 and 19 respectively), years of experimentations.

### Joint regression (FW) and factorial regression analyses

We considered the slopes of the joint regression (slope_FW) and the factorial regression models as measures of individual plasticity (Figs 3B, C). While slope_FW was calculated by regressing the observed phenotypes on the effects of the environment (Supplementary Table S10), the factorial regression slope was calculated with different explicit environmental covariates (Malosetti et al., 2013), which allowed us to assess the contribution of each climatic variable to G×E. Mean temperature (Tmean) was the factor that contributed most to the interaction (Supplementary Table S11), which is consistent with the use of GDD, which takes Tmean into account in its calculation. Moreover, this contribution and that of the GDD were also the most significant when the factorial regression analysis was carried out in calendar days (Supplementary Table S11). Other variables such as photoperiod, photoperiod × GDD and global radiation did not improve the model (Supplementary Table S11). Thus, we calculated the slope using Tmean as covariate (slope_Tmean) (Supplementary Table S11). Notably, the use of Tmean as the environmental index was more efficient to model G×E, as the factorial regression captured a larger variance of G×E (8.5%) than the FW regression (2.8%) (Supplementary Tables S7, S8).

Individuals showed a wide range of slope (slope_FW, slope_Tmean) (Figs 3B, C; Supplementary Table S12). Slopes from both models were highly negatively correlated (R^2^ = 0.91) (Supplementary Fig. S3A). They were also correlated to the overall mean flowering time (R^2^ = 0.67) (Supplementary Figs S3B, C), indicating that early flowering genotypes were on average less stable. Indeed, late genotypes (e.g. ‘H102’) in Southern locations could rank as early or moderate early flowering genotypes in Northern locations, whereas early flowering genotypes (e.g. ‘H056’) in Southern locations could rank as moderately late or late flowering genotypes in Northern Europe (Figs 3B, C).

In addition, the joint regression model estimates a non-linear plasticity parameter, which presumably has a different genetic basis (Kusmec et al., 2017). This parameter is the residual error of the joint regression model (Fig. 3D). Several genotypes, namely ‘H036’, ‘H120’, ‘H105’, ‘H064’ and ‘H065’, presented high residual variances as they displayed a nonlinear response to the environmental gradient (Fig. 3D; Supplementary Table S10). For instance, ‘H036’ was overall ranked as a moderate early flowering genotype but presented a large deviation in FR18, where it was the second earliest genotype.

### Genetic architecture of flowering time

We explored the genetic architecture of flowering time through QTL analysis based on male and female linkage maps with a total of 12196 SNP markers from the Affymetrix® SNP array (Hardigan et al., 2020). The linkage maps were constructed with a total of 6778 and 5418 markers for the female and male linkage maps, respectively. The final number of markers covered the expected 28 linkage groups (LG) for both female and male maps with additional small LG (44 for female and 33 for male linkage maps) (Supplementary Table S13). The lengths of the female and male linkage maps were 2298.5 cM and 1653.1 cM, respectively, with an average distance between markers of 0.7 cM. LG were assigned to one of the seven homoeologous groups (HG) according to the nomenclature of Hardigan et al. (2020) where letters refer to subgenomes (A, B, C, D) and using the Royal Royce genome for LG orientation.

The list of significant QTL and QEI for single and multi-environment models and for plasticity parameters is provided in Table 2. A total of 28 QTL and QEI linked to flowering time were detected and represented on the linkage groups (Fig. 4A). They can be summarized into 10 unique QTL including two QTL on LG7A (Figs 4A, B). Four flowering time QTL were detected only with single-environment means (7A_F, 7A_M) or only with plasticity parameters (4D_F, 6B_M). The multi-environments model allowed the detection of three QEI (2C_M, 3A_M and 6D_M) and six QTL linked to mean flowering times (1B_M, 1C_M, 2C_M, 3A_M, 6A_M and 6D_M) (Fig. 4C). It is noteworthy that the trend of the QTL effect was maintained whatever the environment but its magnitude could vary considerably from one environment to another (Fig. 4D).

**Table 2.**
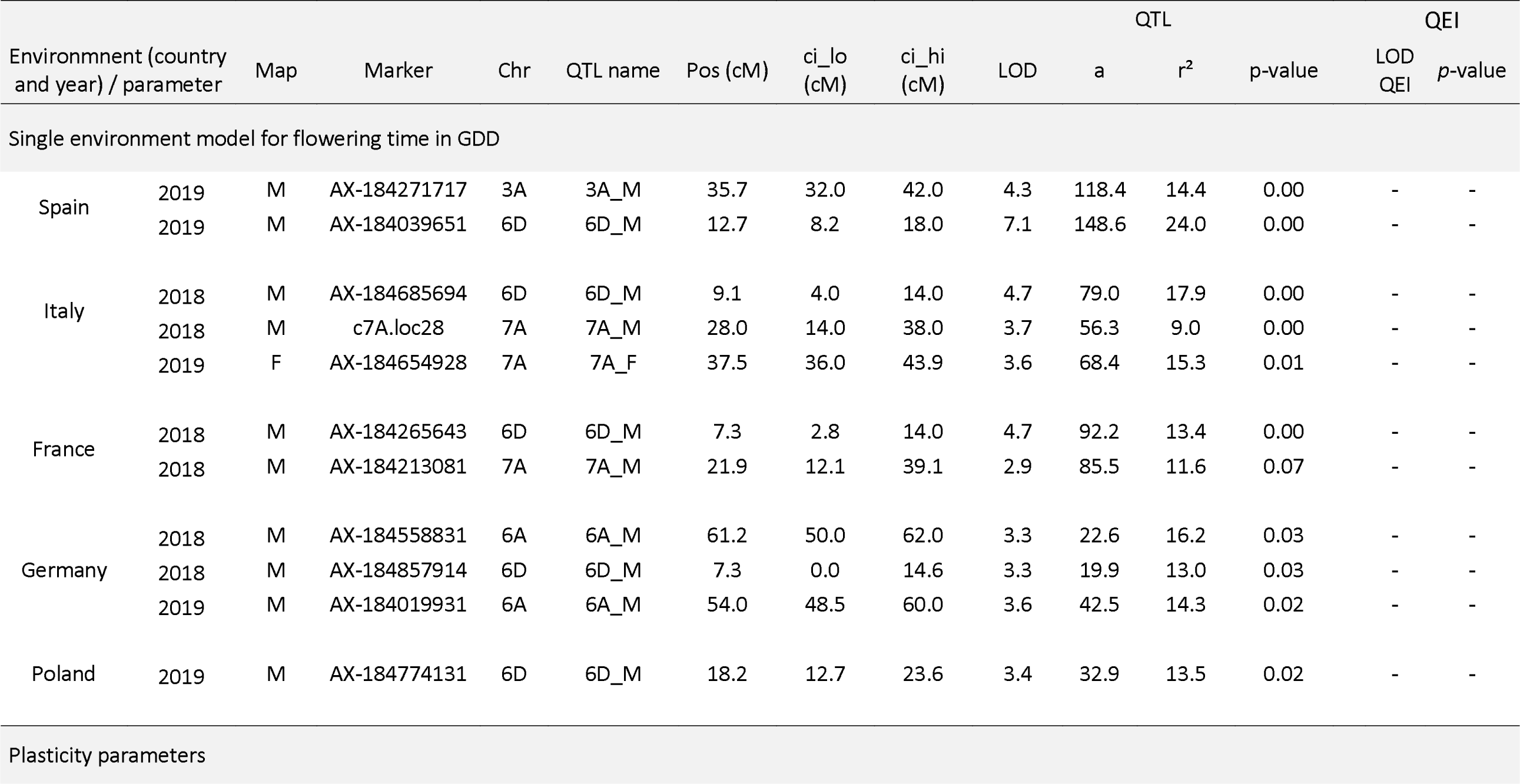
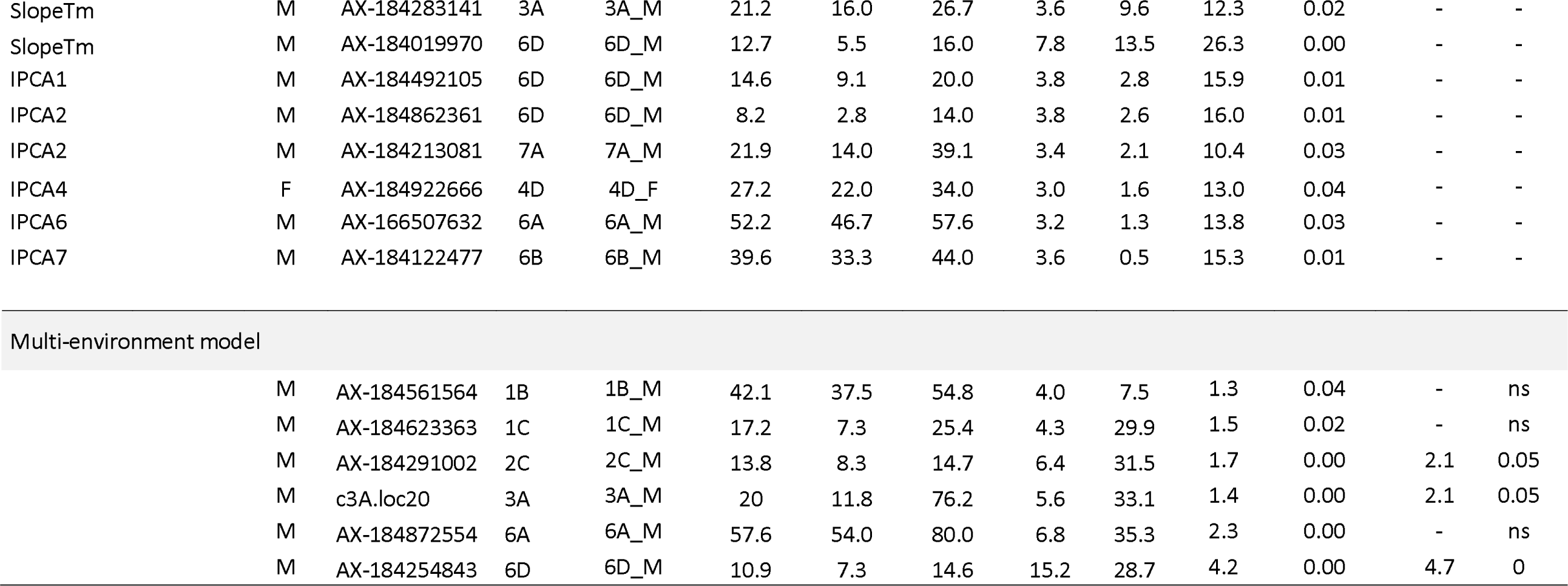
QTL and QEI detected for flowering time in single and multi-environment models and for plasticity parameters. Chr, chromosome; Pos, genetic position in cM; ci_lo, lower genetic position in cM of the Bayesian credible interval; ci_hi, upper genetic position in cM of the Bayesian credible interval; LOD, logarithm of the odds ratio; a, mean phenotypic difference between the two homozygous loci of the QTL; r², percentage of total phenotypic variance explained by the QTL; QEI, QTL-by-Environment Interaction.

**Figure 4.**
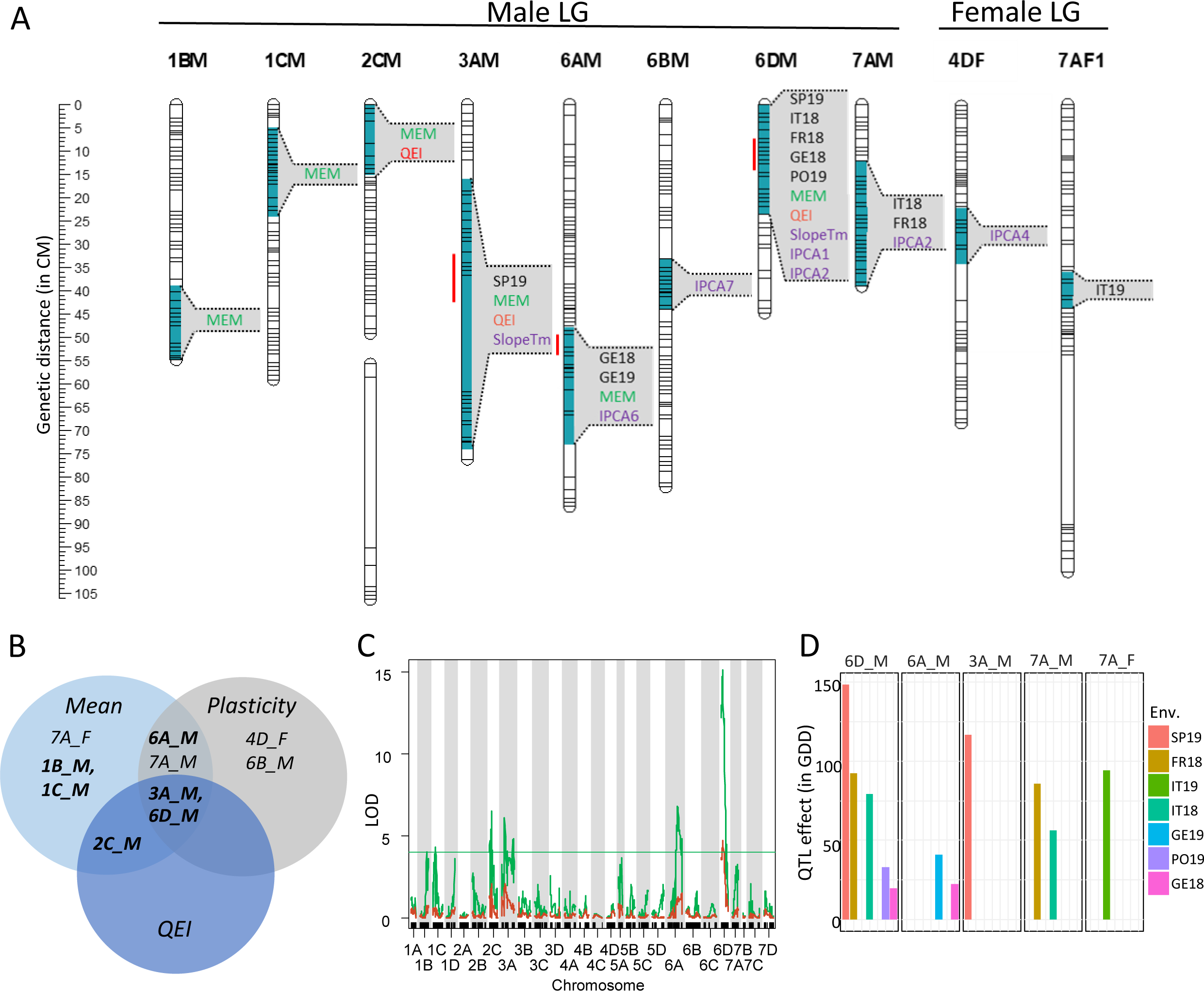
Effects of flowering time QTL and QTL-by-Environment Interactions (QEI). (A) Position of flowering time QTL detected for each environment (in black), for plasticity parameters (slope_Tmean, ICPA1, IPCA2, IPCA4, IPCA6, ICPA7 in purple), and for the overall mean flowering time across the nine environments (in green) obtained using the multi-environment model (MEM). Significant QEI in the MEM are written in red. Linkage groups (LG) are ordered by male and female linkage maps. Red lines, Bayesian credible intervals common to the different QTL detected in 3A_M, 6A_M and 6D_M QTL. (B) Venn diagram of QTL detected for flowering time and plasticity parameters. Mean: QTL detected by environment for the mean flowering times or with the MEM for the overall flowering time; Plasticity parameters: QTL detected for slope_Tmean (slopeTm), ICPA1, IPCA2, IPCA4, IPCA6, ICPA7; QEI: QTL-by-Environment Interactions detected with the MEM. QTL detected for the overall mean with MEM are in bold. (C) LOD scores of QTL and QEI obtained for the multi-environment model: in green the LOD curve for main and interactive effects, in red the LOD curve for the interactive term alone. Thresholds, α = 5%. (D) Variation in QTL effects for flowering time. Only QTL detected by environment are represented (α = 5%). QTL are named according to the LG where they were detected. M and F, male and female linkage maps respectively. SP, Spain, IT, Italy, FR, France, GE, Germany, PL, Poland. 2018 and 2019 (18 and 19 respectively), years of experimentations.

As could be expected from the strong correlations (>0.7) between the overall mean of flowering time and plasticity parameters (Supplementary Fig. S3), we observed co-localizations between them for 3A_M, 6A_M and 6D_M QTL. Of notice, 6A_M QTL was detected in Germany for the two years and for one plasticity parameter (IPCA6). Only two QTL displayed both interaction with the environment and co-localization between the overall mean flowering time and plasticity parameters, being 3A_M and 6D_M QTL. The latest displayed the highest number of co-localizations and the highest LOD values, being detected for five environments and three plasticity parameters (slope_Tmean, IPCA1 and IPCA2) whereas 3A_M QTL was only detected for SP19 and slope_Tmean. The 7A_M QTL was detected only with single-environment means (FR18 and IT18) and one plasticity parameter (IPCA2) (Figs 4A, B, 5A; Table 2).

**Figure 5.**
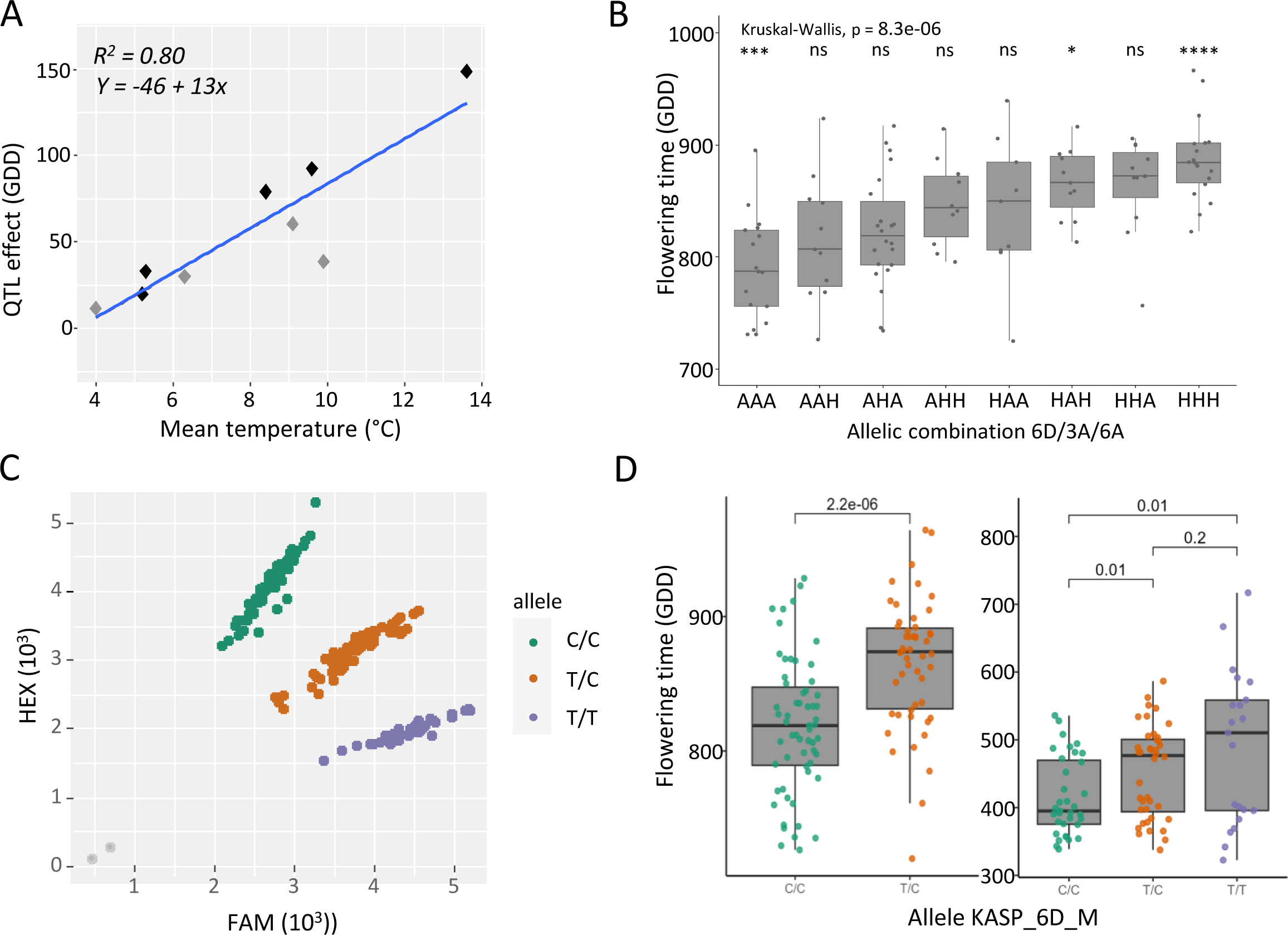
Allelic effects of flowering time QTL. (A) Effect of the 6D_M QTL on flowering time according to mean temperature calculated from the 1^st^ of January until end of flowering. QTL effects: significant (black point) or non-significant (grey point). (B) Effect of alleles of the three major QTL (respectively 6D_M, 3A_M, 6A_M) on flowering time (GDD). Significant pairwise differences levels between allelic classes at the three markers are indicated by stars following a Kruskal-Wallis test (ns, non-significant). (C) Allele Specific PCR (KASP) assay developed on the 6D. The green and purple dots represent the homozygous genotypes (C/C and T/T) and the orange dot represents heterozygous genotypes. The gray dots represent the non-template control. (D) Effect of the KASP_6D marker on flowering time in the ‘Candonga’ x ‘Senga Sengana’ segregating population (left) and in a set of 94 genotypes (right). (D) Allelic effect of the KASP_6D marker on flowering time in the ‘Candonga’ x ‘Senga Sengana’ segregating population (left) and in a set of 94 genotypes (right).

Two results suggest an effect of temperature on the 6D_M QTL: (i) QTL and QEI for the mean flowering times and slopes calculated using the Tmean as covariate (slope_Tmean) are co-located in 6D_M (Fig. 4A) and (ii) the effect decreases from the south to the north of Europe (Fig. 4D). Indeed, we observed that the allelic effect of 6D_M QTL increased linearly (R² = 0.80) with Tmean across environments (Fig. 5A), resulting in a difference of up to 150 GDD (more than six days) in the warmest environment (SP19) but less than 25 GDD (less than one day) in the coldest environment (GE18) (Fig. 4D). Such relation was less clear for the other QEI (2C_M and 3A_M QTL, Supplementary Fig. S4).

We focused more specifically on the effect of alleles associated with the three QTL, 6D_M, 6A_M and 3A_M co-localizing for the overall mean flowering times in the multi-environment model and plasticity parameters (Fig. 5B). The single-marker analysis showed the strongest effect of the allele linked to the 6D_M QTL on flowering time compared with the 3A_M and 6A_M, with a gain of respectively 52, 30 and 35 GDD (Supplementary Fig. S5). The earliest flowering genotypes combined the A alleles for the three markers while the latest flowering genotypes were H for the three markers with an average gain of 97 GDD (Fig. 5B).

### Candidate genes

We identified five candidate genes associated with flowering time within the common Bayesian credible interval of 3A_M, 6A_M and 6D_M QTL (Table 3). We retained candidate genes when they were annotated in both Camarosa va1.0 (Edger et al., 2019) and Royal Royce va1.0 (Hardigan et al., 2021) genomes. In the LG3A_M interval, we identified two candidate genes associated with flowering time: *FaCEN-like* (*CENTRORADIALIS*) and *FaFRI-like* (*FRIGIDA*). In the LG6A_M interval, we identified two flowering-time-related proteins: FaFY and FaFPA. In the LG6D_M QTL interval *FaTFL1* (*TERMINAL FLOWER1*, which belongs to the CENTRORADIALIS/TERMINAL FLOWER 1/SELF-PRUNING (CETS) family, as *FaCEN-like* in LG3A_M, was the most relevant candidate gene.

**Table 3.** Candidate genes identified in the three flowering time QTL common to the overall mean flowering times (multi-environment model) and the plasticity parameters.

| QTL position | Position on Camarosa genome | Position on Royal Royce genome | Candidate gene | ID on Camarosa genome | ID on Royal Royce genome | Corresponding ID on diploid genome | Arabidopsis |
| --- | --- | --- | --- | --- | --- | --- | --- |
| LG3A | 9077738-21022370 | 9867318-22377776 | *CEN-like/ATC* | FxaC_9g27230 | Fxa3Ag102363 | FvH4_3g24700 | AT2G27550 |
|  |  |  | *FRI-like1* | FxaC_9g28150 | Fxa3Ag102288 | FvH4_3g24000 | AT5G16320 |
| LG6A | 7331233-11484453 | 24040354-27479667 | *FY-like* | FxaC_21g17390 | Fxa6Ag103856 | FvH4_6g22190 | AT5G13480 |
|  |  |  | *FPA-like* | FxaC_21g17211 | Fxa6Ag103868 | FvH4_6g40190 | AT4G12640 |
| LG6D | 11903312-15757612 | 9377205-13301702 | *TFL1* | FxaC_24g24110 | Fxa6Dg101555 | FvH4_6g18480 | AT5G03840 |

### Development of a KASP marker for Marker Assisted Selection

We developed a KASP marker linked to the 6D_M QTL (KASP_6D) for further use in Marker-Assisted Selection (MAS) in breeding programs. We analyzed the polymorphism of this marker in our bi-parental population. In addition, we validated its utilization by using a collection of 94 strawberry SF genotypes scored in two successive years in France (Douville). This marker was able to discriminate three genotypes: C/C, T/C, and T/T (Fig. 5C). In the bi-parental population, C/C genotypes required on average 50 fewer heat units (GDD) than T/C genotypes and thus provided earlier flowering (Fig. 5D). Its allelic effect was highest in Spain (a gain of up to 133 heat units (GDD) i.e. almost 8 days) and lowest (a gain of 0-2 days) in Germany and Poland. T/T alleles were exclusively present in the collection of SF genotypes where C/C alleles brought a gain of 73 heat units (GDD) (around 7 days) compared to T/T. The phenotypes of the T/C and T/T genotypes were not significantly different (Fig. 5D). Overall, our results clearly indicate that the introduction of C/C alleles can be effective in the selection of early flowering strawberry varieties, especially in southern Europe.

## Discussion

Flowering time has been extensively characterized in crop species as co-determinant of seed or fruit yield (Jung and Müller, 2009; Eshed and Lippman, 2019). The plasticity of flowering time has been well studied in crops (Li et al., 2018), but the variability of its response to various environmental cues depends on species and/or environmental range (Arnold et al., 2019). The genetic architecture of the plasticity, i.e. the ability of a plant to change its phenotype according to environments, has been investigated in a limited number of crop species (e.g., sorghum, Li et al., 2018; tomato, Diouf et al., 2020; maize, Jin et al., 2023; cherry, Branchereau et al., 2023). To date, no similar effort has been devoted to strawberry.

Here, thanks to a multi-European research program coordinated between five southern and northern European countries representative of the leading strawberry production areas in Europe, we have for the first time dissected the genetic basis of flowering time and its plasticity in relation to environmental cues. To this end, we analyzed the phenotypic response of a segregating population of cultivated strawberry grown in nine contrasting environments using different models. The genetic architecture highlighted both shared and distinct genetic control of flowering time and its plasticity, as well as genetic-based sensitivity to temperature variations.

### The plasticity of flowering time is driven by temperature over a wide range of latitudes

Identifying environmental parameters that have an impact on flowering time is essential to understand the mechanisms underlying its phenotypic plasticity (Mu et al., 2022). Temperature and photoperiod are known as major drivers of this trait (Blackman, 2017). In our study, we showed the predominance of the temperature when modeling flowering time, with thermal time (GDD) having the highest efficiency (85%) when compared to photoperiod (62%) or global radiation (37%) (Table 1). The 15 percent remaining efficiency could be due to differences in cultural techniques among countries (e.g., soilless culture in France or soil culture in Italy), which produce different plant architectures and therefore variations in flowering patterns (Neri et al., 2012).

Plasticity can also be studied through the decomposition of G×E, which reveals the variation in reaction norms between genotypes (Sultan, 1987). We showed here that in strawberry, the G×E variance for flowering time represents a high proportion of the total variance. This result indicates that, when flowering time is considered, strawberry is a very plastic species, more so than suggested for sorghum and cherry (Li et al., 2018; Branchereau et al., 2023). Among the models used for studying G×E, the factorial regression models describe a genetically controlled differential sensitivity to explicit environmental factors (Malosetti et al., 2013). These models, therefore, provide responses as to the climatic drivers of the trait (Lombardi et al., 2022) and environmental indexes that can be used to predict trait performance and inform the design of future studies (Guo et al., 2023). In our study, the factorial regression model used confirmed results from the GDD model by identifying the mean temperature as the dominant climatic factor affecting flowering time, well ahead of photoperiod or global radiation (Supplementary Table S11).

The weaker influence of photoperiod than temperature on flowering time is likely due to the fact that our study was conducted on a population of SF genotypes, the most common type of cultivated strawberry. Photoperiod plays an essential role in floral initiation of strawberry (Heide et al., 2013). In SF strawberry, the dormancy period separates floral initiation from flowering (Gaston et al., 2021) and, therefore, can act as a reset, leading to at least partial independence between these two processes (Krüger et al., 2022). In contrast, in PF strawberry i.e. varieties producing flowers all along their vegetative cycle (Samad et al., 2022), and in forcing cultures with a year-round production system (Yamasaki, 2013), floral initiation is immediately followed by flowering, which may explain why photoperiod can have a greater influence on the timing of flowering.

### Improving the prediction of flowering time in contrasting environments

In the very near future, strawberry production areas will face major variations in both average temperature (Tmean) and maximum temperature (Tmax) as a result of climate change (https://www.ipcc.ch/assessment-report/ar6/). To predict the adaptation of a strawberry variety to various environments using modeling, the parameters of the model must be accurately determined.

To calculate heat accumulation, the GDD model assumes that there is a lower limit temperature (Tb). In strawberry, Tb was imputed arbitrary for blooming at 0°C (Opstad et al., 2011; Bethere et al., 2016) or was calculated for leaf appearance (0°C, Rosa et al., 2011). Our GDD model calculated Tb as -1.7°C. Such negative Tb temperature has been previously described, for example in wheat for leaf appearance (Zartash et al., 2020). Our calculated minimum temperature is thus consistent with the -1.0 to -2.0°C temperatures of the cold rooms used to store plug plants and stop their development until plantation (Lieten et al., 2005).

The GDD model does not predict the maximum temperature (Tmax), the temperature threshold above which additional heat no longer contributes to the calculation of flowering time (Elmendorf and Hollister, 2023). However, knowing the Tmax is necessary to anticipate the high temperatures predicted by climate change models. Using the triangular model, we estimated Tmax at 34°C; this temperature was exceeded only occasionally in our experiments. The Tb and Tmax values found in our study will be incorporated into models to improve flowering time prediction for strawberry, particularly under the hottest conditions (Jochner et al., 2016).

### Integrating the results of the G×E analyses makes it possible to decipher the genetic architecture of flowering time plasticity

In the context of climate change, to overcome the problem of traditionally selected varieties, which are highly efficient but not very plastic, it is becoming increasingly necessary to produce genotypes suitable for multi environments. This can be achieved by taking advantage of phenotypic plasticity in breeding programs (Kusmec et al., 2018; Monforte, 2020). The genetic basis of phenotypic plasticity has been a central research topic in recent decades (Pigliucci, 2005). In this study, by combining the detection of multi-environmental and environment-specific QTL, we have highlighted the differential sensitivity of QTL to environmental changes and the influence of G×E on strawberry flowering phenotype. We observed four QTL displaying co-localization between the mean flowering times (single- and multi-environment models) and plasticity parameters (3A_M, 6A_M, 6D_M and 7A_M). Remarkably, while two of these QTL (3A_M and 6A_M) were identified in a single country (the 3A_M QTL in Spain, the 6A_M QTL in Germany), the 6D_M QTL was detected across multi environments and in the five countries. In the case of the 3A_M, 6A_M and 7A_M QTL, the genetic control of flowering time likely reflects the adaptation of strawberry to local climates (Mitchell-Olds and Schmitt, 2006). The 3A_M QTL could particularly be useful for breeding strawberry varieties adapted to the hotter conditions of Spain and other countries where strawberry production is expanding (for example Morocco and Mexico), whereas the use of the 6A_M QTL could be more relevant in temperate-cold conditions. The 7A_M QTL was a particular case as it was not detected in the multi-environment model and could be linked to specific conditions met in France and Italy in 2018. Remarkably, the sign of the effect of 6D_M QTL was consistent across all environments, meaning that it can be used in breeding programs to create varieties adapted to both northern and southern European climates. However, as its effect on flowering time is higher in southern (subtropical) Europe and lower in northern (temperate-cold) Europe, the use of this QTL for breeding would be more relevant in southern Europe and other countries with similar climates.

Two models have been proposed for the genetic control of phenotypic plasticity (Via et al., 1995): (i) the gene-regulation model, according to which regulatory loci modify the expression of other genes (e.g. structural genes) as a function of the environment, and (ii) the allelic sensitivity model, according to which the alleles underlying the QTL are differentially expressed depending on the environment. These models involve different genetic controls: the gene-regulation model implies that QTL for plasticity parameters are distinct from the mean flowering times QTL, whereas the allelic sensitivity model implies co-localization between them (Gutteling et al., 2007). For four QTL (3A_M, 6A_M, 6D_M and 7A_M), we found co-localization between the mean flowering times QTL and plasticity parameters QTL, which is typically expected for the allelic sensitivity model (Gutteling et al., 2007; Diouf et al., 2020). Most other QTL were either specific to the mean flowering times (e.g. 1B_M, 1C_M and 7A_F) or to one of the plasticity parameters (e.g. 4D_F and6B_M), which is consistent with the gene-regulation hypothesis. Therefore, in the population and environments studied, we found a co-occurrence of the two models in the genetic architecture of flowering time, which is consistent with previous reports on flowering time and other traits in other crop species (Lacaze et al., 2009; Gage et al., 2017; Kusmec et al., 2017; Diouf et al., 2020; Jin et al., 2023).

As cultivated strawberry is an octoploid species (Edger et al., 2019; Gaston et al., 2020), we may hypothesize that strawberry utilizes various homoeoalleles to regulate the timing of flowering depending on the environment, as previously proposed for the control of fruit quality traits (Lerceteau-Köhler et al., 2012). This polyploid plasticity has been postulated to play a considerable role in the evolution of polyploid crop species (Jackson and Chen, 2010). This hypothesis would have far-reaching implications in strawberry breeding as different homoeoalleles of a same gene carried by different chromosomes could contribute to the timing of flowering in changing environmental conditions. However, in strawberry, genomic redundancy does not necessarily translate into greater trait plasticity as previously shown by a study on the influence of polyploidy on the environmental fitness of a series of diploid and polyploid strawberry species (Wei et al., 2019).

### *TFL1*, a likely candidate gene underlying the 6D_M QTL

Several flowering-related genes could be found in the intervals of the three QTL (3A_M, 6A_M, and 6D_M) for which we observed co-localizations between the overall mean flowering times and plasticity parameters. Among these, 3A_M and 6D_M QTL are both sensitive to temperature. The *A. thaliana* homologous of *FaCEN-like* candidate gene underlying the 3A_M QTL has been shown to prolong vegetative growth and consequently delays flowering (Amaya et al., 1999). The *A. thaliana* homolog of the *FaFRI-like* candidate gene that is also found in the 3A_M QTL encodes a transcription factor that positively regulates the expression of *FLOWERING LOCUS C (FLC)* and plays a role in the regulation of natural variation in flowering time in *A. thaliana* (Michaels et al., 2004). Interestingly, Zhu et al. (2021) suggested that a temperature-controlled nuclear condensation mechanism modulates the *FRI* activation of *FLC* transcription, thus contributing to the repression of flowering. The *FaFY* and *FaFPA* candidate genes found in the 6A_M QTL are both known to play a role in *A. thaliana* in the regulation of flowering time in the autonomous flowering pathway by acting on FLC (Koornneef et al., 1991, Cheng et al., 2017).

The *FaTFL1* candidate gene underlying the LG6D_M QTL belongs to the CENTRORADIALIS/TERMINAL FLOWER 1/SELF-PRUNING (CETS) family which plays a pivotal role in either activating or repressing flowering (Wickland and Hanzawa, 2015). The role of TFL1 proteins as major floral repressors is conserved in several species, including tomato (Pnueli et al., 1998) and strawberry (Iwata et al., 2012; Nakano et al., 2015; Koskela et al., 2016). To date, in the diploid strawberry *Fragaria vesca*, the only study on the genetic architecture of flowering time (Samad et al., 2017) was unable to highlight the role of *FvTFL1* in the variation of this trait, as all the genotypes studied were *Fvtfl1* PF mutants. However, *FvTFL1* probably plays a role in regulating the flowering time in *F. vesca*, as its expression is regulated by temperature, being down-regulated at cool temperatures (<13°C) and up-regulated at higher temperatures (23°C); moreover, these features are independent of photoperiod (Rantanen et al., 2015). In cultivated strawberry, *FaTFL1* sensitivity to temperature has been shown to vary according to the genotype: in ‘Elsanta’, *FaTFL1* expression was increased from 9°C to 21°C whereas the temperature had no effect in ‘Glima’ (Koskela et al., 2016). We can assume that one of the two *FaTFL1* homoeoalleles located in the LG6D_M QTL is expressed more in subtropical conditions where temperatures are higher; consequently, this *FaTFL1* allele would delay flowering more significantly in subtropical conditions than in cold temperature conditions. Future studies in controlled conditions will test this hypothesis, for example by carrying out a RNAseq analysis of plant tissues (leaf and bud) collected from genotypes carrying different *FaTFL1* alleles and grown at different temperatures.

## Conclusion

In the context of climate change, it is necessary to uncover the genetic architecture of the plasticity of complex traits in cultivated species. Here, in a concerted European effort, we studied flowering time, a trait that is highly sensitive to the environment, and showed that temperature is the most significant driver of this trait in cultivated strawberry. The detection of several QTL and the identification of underlying candidate genes associated with flowering time plasticity will help to better understand the molecular mechanisms responsible for variations in flowering time and select superior strawberry varieties that are well suited to changing environmental conditions. To this end, we designed the breeder-friendly genetic marker KASP_6D for a major temperature-sensitive QTL, which will accelerate MAS selection for flowering time in cultivated strawberry. Our study will have far-reaching implications for the selection of new strawberry varieties adapted not only to the wide differences in climatic conditions found in Europe, but also to countries with tropical/sub-tropical climates where strawberry production is expanding rapidly.

## Acknowledgements

The authors thank Philippe Chartier, Cédric Duranton and Christian Gauthier (Invenio), Luis Miranda and José A. Gómez-Mora (IFAPA) for strawberry culture and phenotyping. The authors thank Drs. Isidore Diouf and Iñaki Garcia de Cortazar Atauri for fruitful discussions.

## Authors contributions

KO chose the cross ‘Candonga’ × ‘Senga Sengana’ and produced the 1^st^ generation of individuals; BD and members of the GoodBerry project conceived and designed the experiments. AleP and BD analyzed and interpreted the data; BD and AleP wrote the manuscript and CR contributed to the writing of the manuscript. All authors discussed the results and commented on the manuscript.

## Conflict of interest

No conflict of interest declared.

## Funding

The project was funded by European Union’s Horizon 2020 research and innovation program (BreedingValue project N° 101000747). AleP PhD has been supported by Invenio and the Association Nationale de la Recherche et la Technologie (ANRT) (Cifre agreement).

## Data availability

Data will be made available on request.

## Supplementary data

Supplementary Figures

Supplementary Figure S1. Culture workflow of the European Goodberry project.

Supplementary Figure S2. Biplots of the first three components of AMMI analysis of flowering time.

Supplementary Figure S3. Relationship between GDD, slope_FW and slope_Tmean.

Supplementary Figure S4. Relationship between the QTL effect expressed in GDD and the temperature.

Supplementary Figure S5. Single marker analysis showing the effect of the Affymetrix® allele linked to the flowering time QTL.

**Supplementary Tables**

Supplementary Table S1. Name and sequence of primers used for the validation of the 6D_M QTL. Tm, annealing temperature.

Supplementary Table S2. Descriptions of the nine environments.

Supplementary Table S3. Spearman phenotypic correlations for flowering time between environments.

Supplementary Table S4. ANOVA with mixed model, where genotype and genotype-environment interaction (G×E) are random effects.

Supplementary Table S5. Broad-sense heritabilities (H2) of flowering time by environment, country and whole-design level.

Supplementary Table S6. ANOVA for Additive Main Effects and Multiplicative Interaction (AMMI) model applied to flowering time (GDD) in the segregating population ‘Candonga’ x ‘Senga Sengana’ under nine environments.

Supplementary Table S7. ANOVA using joint regression (FW) model on flowering time expressed in GDD.

Supplementary Table S8. ANOVA using factorial regression model on flowering time expressed in GDD or in calendar day.

Supplementary Table S9. AMMI Stability Value (ASV) calculated on the first two ICPA for flowering time (GDD).

Supplementary Table S10. Linear (slope_FW) and non-linear (residual variance, VAR_FW) plasticity.

Supplementary Table S11. Factorial regression models incorporating environmental information as covariate.

Supplementary Table S12. Linear (slope_Tmean) plasticity provided by the factorial regression model on Tmean.

Supplementary Table S13. Summary of the linkage maps of the segregating population ‘Candonga’ x ‘Senga Sengana’.

## Notes

### Competing Interest Statement

The authors have declared no competing interest.

