## Supplemental Figures S1-S5 and Tables S1-S13 for "Strawberry phenotypic plasticity in flowering time is driven by interaction between genetic loci and temperature"

**Title**

**Abstract**

The flowering time, which determines when the fruits or seeds can be harvested, is known to be sensitive to plasticity, i.e. the ability of a genotype to display different phenotypes in response to environmental variations. In the context of climate change, strawberry breeding can take advantage of phenotypic plasticity to create high-performing varieties adapted either to local conditions or to a wide range of climates. To decipher how the environment affects the genetic architecture of flowering time in cultivated strawberry (*Fragaria ×ananassa*) and modify its QTL effects, we used a bi-parental segregating population grown for two years at widely divergent latitudes (5 European countries) and combined climatic variables with genomic data (Affymetrix® SNP array). We detected 10 unique flowering time QTL and demonstrated that temperature modulates the effect of plasticity-related QTL.

We propose candidate genes for the three main plasticity QTL, including *FaTFL1* which is the most relevant candidate in the interval of the major temperature-sensitive QTL (6D_M). We further designed and validated a genetic marker for the 6D_M QTL which offers great potential for breeding programs, for example for selecting of early-flowering strawberry varieties well adapted to different environmental conditions.

Supplementary Table S1. Name and sequence of primers used for the validation of the 6D_M QTL. Tm, annealing temperature.

| Primer name | Type Queue | Sequence 5´-3' | Affymetrix marker | Mutation | Tm (°C) | Product (bp) |
|---|---|---|---|---|---|---|
| KASP_6DM-com-Fw-01 | None | TCATTTGACCACAAGTTACCGATG | AX-184201950 | -- | 59.3 | 81 |
| KASP_6DM-spe-Rev-02 | Fam | gaaggtgaccaagttcatgctGACATAGTCATTGAATACAGTGATAATCCAATG | AX-184201950 | C | 63.3 | |
| KASP_6DM-spe-Rev-01 | Vic(HEX) | gaaggtcggagtcaacggattAGACATAGTCATTGAATACAGTGATAATCCAATA | AX-184201950 | T | 62.2 | |

Supplementary Table S2. Descriptions of the nine environments. Spain, SP; Italy, IT; France, FR; Germany, GE; Poland, PL. Years of plantation 2018 (18) and 2019 (19). Window: Number of days from the 1st of January to the end of flowering. Tm, Tmean; Tmin, minimum temperature; Tmax, maximum temperature; Amp.th, day-night amplitude; Glob.Rad., global radiation. GDD (sum of Growing Degree Days), and photoperiod were calculated according to the window.

| Env. | Location | Country | Year of planting | Cultivation system | Latitude | End of flowering (julian date) | Window | Tm (°C) | Tmin (°C) | Tmax (°C) | Amp.Th (°C) | Glob.Rad (kw/m²) | GDD (°C) (tb = -1.7°C) | Photoperiod (hours) |
|---|---|---|---|---|---|---|---|---|---|---|---|---|---|---|
| SP19 | Huelva | Spain | 2018 | Soil | 37°24'N | 05/04/2019 | 95 | 13.6 | 7.1 | 24.7 | 17.6 | 1119 | 1795.5 | 1045 |
| IT18 | Agugliano | Italy | 2017 | Soil | 43°32'N | 19/04/2018 | 109 | 8.4 | 1.6 | 17.7 | 16.1 | 855 | 2060.1 | 1206 |
| IT19 | Agugliano | Italy | 2018 | Soil | 43°32'N | 28/04/2019 | 118 | 9.1 | -0.2 | 21.7 | 21.9 | 1171 | 2230.2 | 1329 |
| FR18 | Douville | France | 2017 | Soilless | 45°59'N | 23/04/2018 | 113 | 9.6 | 6 | 14.1 | 8.1 | 2321 | 2135.7 | 1255 |
| FR19 | Douville | France | 2018 | Soilless | 45°59'N | 15/04/2019 | 105 | 9.9 | 4 | 18.7 | 14.7 | 2654 | 1984.5 | 1144 |
| GE18 | Dresden | Germany | 2017 | Soil | 51°05'N | 09/05/2018 | 129 | 5.2 | 1 | 9.1 | 8.1 | 986 | 2438.1 | 1465 |
| GE19 | Dresden | Germany | 2018 | Soil | 51°05'N | 20/05/2019 | 140 | 6.3 | 2.2 | 10.3 | 8.1 | 1021 | 2646 | 1650 |
| PL18 | Skierniewice | Poland | 2017 | Soil | 51°95'N | 12/05/2018 | 132 | 4 | -0.7 | 8.7 | 9.3 | 946 | 2494.8 | 1507 |
| PL19 | Skierniewice | Poland | 2018 | Soil | 51°95'N | 24/05/2019 | 144 | 5.3 | 0.9 | 10 | 9.1 | 939 | 2721.6 | 1695 |

Supplementary Table S3. Spearman phenotypic correlations for flowering time between environments. Correlation (r) and *p* values.

| row | column | Spearman cor. (r) | *p* |
|---|---|---|---|
| IT18 | FR18 | 0.59 | 0 |
| SP19 | PL19 | 0.57 | 0 |
| IT18 | GE18 | 0.53 | 0 |
| IT19 | FR19 | 0.47 | 0 |
| FR19 | GE18 | 0.46 | 0 |
| FR18 | GE18 | 0.45 | 0 |
| IT18 | IT19 | 0.46 | 0 |
| IT19 | FR18 | 0.45 | 0 |
| IT19 | GE18 | 0.45 | 0 |
| FR18 | FR19 | 0.41 | 0 |
| FR18 | GE19 | 0.4 | 0 |
| GE18 | GE19 | 0.36 | 0.0001 |
| IT18 | FR19 | 0.36 | 0.0001 |
| IT18 | GE19 | 0.35 | 0.0002 |
| IT19 | PL19 | 0.36 | 0.0002 |
| GE18 | PL19 | 0.34 | 0.0004 |
| SP19 | FR19 | 0.33 | 0.0004 |
| FR19 | PL19 | 0.33 | 0.0004 |
| SP19 | PL18 | 0.32 | 0.0005 |
| SP19 | IT19 | 0.34 | 0.0005 |
| FR18 | PL19 | 0.32 | 0.0007 |
| SP19 | FR18 | 0.32 | 0.0007 |
| FR19 | GE19 | 0.32 | 0.0007 |
| PL18 | PL19 | 0.31 | 0.001 |
| SP19 | IT18 | 0.28 | 0.0032 |
| FR18 | PL18 | 0.27 | 0.0045 |
| IT18 | PL19 | 0.26 | 0.0056 |
| IT19 | GE19 | 0.26 | 0.008 |
| SP19 | GE19 | 0.23 | 0.0173 |
| SP19 | GE18 | 0.2 | 0.0328 |
| FR19 | PL18 | 0.18 | 0.0521 |
| GE18 | PL18 | 0.18 | 0.0653 |
| GE19 | PL19 | 0.15 | 0.1127 |
| IT19 | PL18 | 0.15 | 0.1331 |
| GE19 | PL18 | 0.09 | 0.3331 |
| IT18 | PL18 | 0.09 | 0.3464 |

Supplementary Table S4. ANOVA with mixed model, where genotype and genotype-environment interaction (G×E) are random effects. Analyses were performed with calendar days and GDD and at the whole design (nine environments) and country levels. GxL, genotype by Location; GxY, Genotype by Year interaction.

**Calendar days**

**All sites**

| Whole design Source | npar | logLik | AIC | LRT | Df | Pr(>Chisq) | Signif | Variance | Total Variance (%) |
|---|---|---|---|---|---|---|---|---|---|
| Intercept | 12 | -22173 | 44371 | | | - | | - | - |
| (1 \| geno) | 11 | -22254 | 44529 | 160.9 | 1 | < 2.2e-16 | *** | 12 | **22.1** |
| (1 \| geno:env) | 11 | -24862 | 49747 | 5378 | 1 | < 2.2e-16 | *** | 28.5 | **52.6** |

By country

| GxYxL Source | npar | logLik | AIC | LRT | Df | Pr(>Chisq) | Signif | Variance | Total Variance (%) |
|---|---|---|---|---|---|---|---|---|---|
| Intercept | 10 | -23886 | 47791 | - | - | - | | - | - |
| (1 \| geno) | 9 | -23902 | 47823 | 33.61 | 1 | 6.73E-09 | *** | 9.6 | **15.9** |
| (1 \| geno:year) | 9 | -24120 | 48257 | 467.78 | 1 | < 2.2e-16 | *** | 5.5 | **9.1** |
| (1 \| geno:location) | 9 | -24999 | 50016 | 2226.9 | 1 | < 2.2e-16 | *** | 20.3 | **33.5** |

**GDD**

**All sites**

| Whole design Source | npar | logLik | AIC | LRT | Df | Pr(>Chisq) | Signif | Variance | Total Variance (%) |
|---|---|---|---|---|---|---|---|---|---|
| Intercept | 12 | -42595 | 85215 | | - | - | | - | - |
| (1 \| geno) | 11 | -42670 | 85362 | 149.8 | 1 | < 2.2e-16 | *** | 2271.2 | **20.7** |
| (1 \| geno:env) | 11 | -45068 | 90158 | 4945.5 | 1 | < 2.2e-16 | *** | 5694.2 | **51.9** |

By country

| GxYxL **Source** | **npar** | **logLik** | **AIC** | **LRT** | **Df** | **Pr(>Chisq)** | **Signif** | **Variance** | **Total Variance (%)** |
|---|---|---|---|---|---|---|---|---|---|
| Intercept | 10 | -43716 | 87452 | | | | | - | - |
| (1 \| geno) | 9 | -43734 | 87485 | 35.12 | 1 | 3.09E-09 | *** | 1989.8 | **16** |
| (1 \| geno:year) | 9 | -43927 | 87872 | 421.88 | 1 | < 2.2e-16 | *** | 938.8 | **7.5** |
| (1 \| geno:location) | 9 | -45025 | 90068 | 2617.3 | 1 | < 2.2e-16 | *** | 4914 | **39.5** |
| Signif. codes: 0 ‘***’ 0.001 ‘**’ 0.01 ‘*’ 0.05 ‘.’ 0.1 ‘ ’ 1 | | | | | | | | | |

Supplementary Table S5. Broad-sense heritabilities (H2) of flowering time by environment, country and whole-design level.

| | Calendar day | | | GDD | | |
|---|---|---|---|---|---|---|
| PL18 | 0.9134 | 0.3961 | 0.9882 | 0.9164 | 0.3409 | 0.8228 |
| PL19 | 0.9253 | | | 0.9197 | | |
| GE18 | 0.9115 | 0.587 | | 0.9232 | 0.3244 | |
| GE19 | 0.9838 | | | 0.9845 | | |
| FR18 | 0.9762 | 0.9137 | | 0.9744 | 0.8825 | |
| FR19 | 0.9738 | | | 0.971 | | |
| IT18 | 0.982 | 0.9212 | | 0.9781 | 0.8939 | |
| IT19 | 0.9611 | | | 0.9594 | | |
| SP19 | 0.9492 | 0.9492 | | 0.9516 | 0.9516 | |

Supplementary Table S6. ANOVA for Additive Main Effects and Multiplicative Interaction (AMMI) model applied to flowering time (GDD) in the segregating population ‘Candonga’ x ‘Senga Sengana’ under nine environments.

GDD

| All env. | Df | Sum Sq | Mean Sq | F value | Pr(>F) | % GxE | Cumulated % GxE |
|---|---|---|---|---|---|---|---|
| ENV | 8 | 59 557 636.45 | 7 444 704.56 | 2 475.36 | 0 | | |
| GEN | 110 | 21 886 703.35 | 198 970.03 | 66.16 | 0 | | |
| ENV:GEN | 868 | 39 901 364.39 | 45 969.31 | 15.28 | 0 | | |
| IPCA1 | 117 | 8 084 751.62 | 69 100.44 | 22.97 | 0 | 23.93 | 23.93 |
| IPCA2 | 115 | 6 170 066.53 | 53 652.75 | 17.83 | 0 | 18.26 | 42.19 |
| IPCA3 | 113 | 4 605 027.23 | 40 752.45 | 13.54 | 0 | 13.63 | 55.82 |
| IPCA4 | 111 | 3 910 573.43 | 35 230.39 | 11.71 | 0 | 11.57 | 67.39 |
| IPCA5 | 109 | 3 205 683.64 | 29 409.94 | 9.77 | 0 | 9.49 | 76.88 |
| IPCA6 | 107 | 2 490 701.94 | 23 277.59 | 7.74 | 0 | 7.37 | 84.25 |
| IPCA7 | 105 | 2 093 270.92 | 19 935.91 | 6.63 | 0 | 6.2 | 90.45 |
| IPCA8 | 103 | 1 792 957.90 | 17 407.36 | 5.79 | 0 | 5.31 | 95.75 |
| IPCA9 | 101 | 1 434 834.54 | 14 206.28 | 4.72 | 0 | 4.25 | 100 |
| Residuals | 6 610.00 | 19 879 722.86 | 3 007.52 | NA | NA | | |

Supplementary Table S7. ANOVA using joint regression model on flowering time expressed in GDD.

| Factor | df | Sum Sq | Mean Sq | F value | Pr(>F) | PropSSq |
|---|---|---|---|---|---|---|
| *GDD* | | | | | | |
| Genotype | 110 | 2.91E+06 | 2.64E+04 | 4.11 | 3.24E-31 | 18.3 |
| Env | 8 | 7.69E+06 | 9.61E+05 | 149.6 | 2.48E-150 | 48.3 |
| Genotype:score_Env | 110 | 4.44E+05 | 4.03E+03 | 0.63 | 9.99E-01 | 2.8 |
| Residuals | 758 | 4.87E+06 | 6.43E+03 | NA | NA | 30.6 |

Supplementary Table S8. ANOVA using factorial regression model on flowering time expressed in GDD or in calendar day.

| Factor | Df | Sum Sq | Mean Sq | F value | Pr(>F) | PropSSq |
|---|---|---|---|---|---|---|
| *GDD* | | | | | | |
| Genotype | 110 | 2 906 839.39 | 26 425.81 | 5.06 | 6.82E-42 | 18.3 |
| Env | 8 | 7 689 812.61 | 961 226.58 | 184 | 6.45E-172 | 48.3 |
| Genotype:Tm | 110 | 1 354 288.06 | 12 311.71 | 2.36 | 1.81E-11 | 8.5 |
| Residuals | 758 | 3 959 748.05 | 5 223.94 | NA | NA | 24.9 |
| *Calendar day* | | | | | | |
| Genotype | 110 | 15 825.22 | 143.87 | 5.15 | 7.20E-43 | 2.9 |
| Env | 8 | 500 483.34 | 62 560.42 | 2238 | 0.00E+00 | 92.2 |
| Genotype:Tm | 110 | 5 197.85 | 47.25 | 1.69 | 4.56E-05 | 1 |
| Residuals | 758 | 21 189.04 | 27.95 | NA | NA | 3.9 |

Supplementary Table S9. AMMI Stability Value (ASV) calculated on the first two ICPA for flowering time (GDD).

| Geno | ASV | Geno | ASV | Geno | ASV |
|---|---|---|---|---|---|
| Candonga | 0.13 | H043 | 0.28 | H085 | 0.25 |
| H001 | 0.69 | H045 | 0.87 | H086 | 0.37 |
| H002 | 0.17 | H046 | 0.82 | H087 | 0.79 |
| H003 | 0.27 | H047 | 0.55 | H088 | 0.40 |
| H004 | 0.60 | H049 | 0.50 | H089 | 0.34 |
| H005 | 0.80 | H050 | 1.07 | H090 | 0.20 |
| H006 | 0.80 | H051 | 0.83 | H091 | 0.06 |
| H007 | 0.33 | H052 | 0.68 | H093 | 0.08 |
| H008 | 0.62 | H053 | 0.54 | H094 | 0.33 |
| H009 | 0.21 | H054 | 0.44 | H095 | 0.59 |
| H010 | 0.16 | H055 | 0.54 | H096 | 0.41 |
| H011 | 0.65 | H056 | 0.25 | H097 | 0.35 |
| H012 | 0.62 | H057 | 0.19 | H098 | 0.11 |
| H013 | 0.28 | H058 | 0.20 | H101 | 0.35 |
| H014 | 0.43 | H059 | 0.64 | H102 | 0.12 |
| H015 | 0.18 | H060 | 0.59 | H103 | 0.35 |
| H016 | 0.11 | H062 | 0.14 | H104 | 0.07 |
| H017 | 0.54 | H063 | 0.67 | H105 | 0.28 |
| H019 | 0.46 | H064 | 0.22 | H106 | 0.49 |
| H020 | 0.74 | H065 | 0.41 | H107 | 0.20 |
| H021 | 0.60 | H066 | 0.21 | H110 | 0.45 |
| H022 | 0.56 | H067 | 0.46 | H112 | 0.58 |
| H023 | 0.66 | H068 | 0.65 | H113 | 1.03 |
| H024 | 1.09 | H070 | 0.55 | H114 | 1.12 |
| H027 | 1.30 | H071 | 0.27 | H115 | 0.87 |
| H028 | 0.68 | H072 | 0.77 | H116 | 0.21 |
| H029 | 0.24 | H073 | 1.20 | H117 | 0.34 |
| H030 | 0.61 | H074 | 0.43 | H118 | 0.35 |
| H031 | 0.39 | H075 | 0.24 | H119 | 0.89 |
| H032 | 0.65 | H076 | 0.38 | H120 | 0.33 |
| H033 | 0.59 | H077 | 0.07 | H121 | 0.99 |
| H035 | 0.25 | H078 | 0.35 | H122 | 1.17 |
| H036 | 0.52 | H079 | 0.55 | H123 | 0.90 |
| H037 | 0.49 | H080 | 0.11 | H124 | 0.50 |
| H039 | 0.49 | H081 | 0.18 | H125 | 0.88 |
| H040 | 0.20 | H083 | 0.92 | H126 | 0.43 |
| H042 | 0.40 | H084 | 0.16 | SengaSengana | 0.65 |

Supplementary Table S10. Linear (slope_FW) and non-linear (residual variance, VAR_FW) plasticity. Values were provided by the Finlay Wilkinson (FW) model (Kusmec et al., 2017) for each individual of the segregating population 'Candonga' x 'Senga Sengana' evaluated under nine environments. FW model was applied on flowering time (GDD).

| Geno | Slope_FW | VAR_FW |
|---|---|---|
| H096 | 0.74 | 17.46 |
| H102 | 0.76 | 43.73 |
| H116 | 0.81 | 36.52 |
| H080 | 0.83 | 19.84 |
| H047 | 0.84 | 21.25 |
| H007 | 0.86 | 32.27 |
| H046 | 0.87 | 33.63 |
| H006 | 0.88 | 25.93 |
| H015 | 0.88 | 11.03 |
| H051 | 0.88 | 41.19 |
| H023 | 0.89 | 1.71 |
| H049 | 0.89 | 2.69 |
| H039 | 0.89 | 35.65 |
| H031 | 0.9 | 27.97 |
| H043 | 0.91 | 21.97 |
| H009 | 0.91 | 3.96 |
| H123 | 0.91 | 32.99 |
| H125 | 0.91 | 18.68 |
| H019 | 0.92 | 18.17 |
| H094 | 0.92 | 9.21 |
| H090 | 0.92 | 7.79 |
| H028 | 0.92 | 11.38 |
| H002 | 0.92 | 6.72 |
| H012 | 0.92 | 18.23 |
| H081 | 0.93 | 10.91 |

| Geno | Slope_FW | VAR_FW |
|---|---|---|
| H004 | 0.95 | 6.35 |
| H114 | 0.95 | 14.39 |
| H083 | 0.96 | 25.3 |
| H063 | 0.96 | 32.84 |
| H089 | 0.96 | 6.23 |
| H060 | 0.96 | 6.31 |
| H064 | 0.96 | 64.2 |
| H115 | 0.96 | 5.94 |
| H027 | 0.97 | 25.81 |
| H122 | 0.97 | 10.92 |
| H093 | 0.97 | 35.66 |
| H014 | 0.97 | 6.24 |
| H097 | 0.98 | 53.38 |
| Candonga | 0.98 | 37.84 |
| H032 | 0.98 | 45.51 |
| H003 | 0.98 | 46.46 |
| H101 | 0.98 | 9.81 |
| H104 | 0.98 | 38.12 |
| H073 | 0.99 | 28.63 |
| H068 | 0.99 | 8.58 |
| H024 | 0.99 | 35.55 |
| H086 | 1 | 58.14 |
| H071 | 1 | 19.34 |
| H070 | 1 | 22.83 |
| H021 | 1.01 | 30.62 |

| Geno | Slope_FW | VAR_FW |
|---|---|---|
| H005 | 1.03 | 27.65 |
| H072 | 1.04 | 14.13 |
| H078 | 1.04 | 28.81 |
| H085 | 1.04 | 12.68 |
| H042 | 1.05 | 15.71 |
| H022 | 1.05 | 31.35 |
| H052 | 1.05 | 17.54 |
| H013 | 1.05 | 27.77 |
| H066 | 1.06 | 12.9 |
| H110 | 1.06 | 12.71 |
| H077 | 1.06 | 17.31 |
| H059 | 1.06 | 19.13 |
| H050 | 1.06 | 2.6 |
| H040 | 1.07 | 14.75 |
| H065 | 1.07 | 9.59 |
| H055 | 1.07 | 19.14 |
| H124 | 1.08 | 13.18 |
| H008 | 1.09 | 17.07 |
| H105 | 1.09 | 67.22 |
| H091 | 1.09 | 23.62 |
| H113 | 1.11 | 7.3 |
| H054 | 1.11 | 7.53 |
| H037 | 1.12 | 4.1 |
| H118 | 1.13 | 23.72 |
| H121 | 1.13 | 24.24 |

| | | |
|---|---|---|
| H075 | 0.93 | 36.01 |
| H029 | 0.93 | 38.75 |
| H103 | 0.94 | 13.97 |
| H045 | 0.94 | 27.21 |
| H084 | 0.94 | 19.79 |
| H030 | 0.94 | 41.45 |
| H001 | 0.94 | 58.82 |
| H095 | 0.94 | 16.38 |
| H079 | 0.95 | 36.03 |
| H058 | 0.95 | 26.25 |
| H011 | 0.95 | 11.67 |
| H117 | 0.95 | 46.05 |

| | | |
|---|---|---|
| H106 | 1.01 | 27.66 |
| H088 | 1.01 | 31.74 |
| H053 | 1.01 | 10.95 |
| H036 | 1.01 | 83.03 |
| H098 | 1.01 | 5.08 |
| H020 | 1.01 | 3.62 |
| H112 | 1.02 | 32.98 |
| SengaSengana | 1.02 | 39.46 |
| H107 | 1.03 | 13.64 |
| H087 | 1.03 | 45.07 |
| H074 | 1.03 | 12.11 |
| H076 | 1.03 | 14.28 |

| | | |
|---|---|---|
| H016 | 1.13 | 37.64 |
| H010 | 1.15 | 38.34 |
| H035 | 1.15 | 13.05 |
| H067 | 1.16 | 14.9 |
| H017 | 1.19 | 32.33 |
| H126 | 1.19 | 31.32 |
| H033 | 1.19 | 49.36 |
| H062 | 1.2 | 14.14 |
| H057 | 1.21 | 15.61 |
| H120 | 1.22 | 70.34 |
| H056 | 1.23 | 39.58 |
| H119 | 1.25 | 31.23 |
| MEAN | 1 | 24.66 |

Supplementary Table S11. Factorial regression models incorporating environmental information as covariate. Tm, Tmean; GDD, Growing Degree Day; Tmax, maximum temperature; Tmin, minimum temperature; Amp.th, temperature difference between Tmin and Tmax; Glob.Rad, Global Radiation; PhotoP, Photoperiod; TmxPhotoP, photothermal time. Analyses were performed on flowering time (GDD or calendar days). SSQ, Sum of SQuare.

GDD

| Covariate | Tm | GDD | Tmax | Tmin | Amp.Th | Glob.Rad | PhotoP. | TmxPhotoP. |
|---|---|---|---|---|---|---|---|---|
| SSQ | 1 354 288.06 | 1 328 464.89 | 1 243 330.44 | 1 197 075.85 | 982 535.52 | 645 030.14 | 970 808.57 | 1 272 611.95 |
| Pvalue | 1.81E-11 | 7.64E-11 | 6.88E-09 | 6.71E-08 | 0.000477 | 0.618928891 | 0.0007 | 1.53E-09 |

Calendar days

| Covariate | Tm | GDD | Tmax | Tmin | Amp.Th | Glob.Rad | PhotoP. | TmxPhotoP. |
|---|---|---|---|---|---|---|---|---|
| SSQ | 5 031.60 | 5 197.85 | 4 836.18 | 4 767.78 | 4 530.74 | 3 238.71 | 3 911.43 | 4 773.89 |
| Pvalue | 1.61E-04 | 4.56E-05 | 6.39E-04 | 1.01E-03 | 4.44E-03 | 5.85E-01 | 9.30E-02 | 9.71E-04 |

Supplementary Table S12. Linear (slope_Tmean) plasticity provided by the factorial regression model on Tmean.

| Geno | Slope_Tmean | Geno | Slope_Tmean | Geno | Slope_Tmean |
|---|---|---|---|---|---|
| H056 | -33.47 | H112 | -9.02 | H046 | 3.81 |
| H119 | -31.41 | H078 | -7.64 | H123 | 4.17 |
| H057 | -31.34 | H088 | -7.35 | H081 | 5.14 |
| H033 | -27.54 | H098 | -7.23 | H114 | 5.51 |
| H118 | -25.50 | H104 | -7.00 | H024 | 5.75 |
| H017 | -23.08 | H020 | -6.79 | H090 | 5.81 |
| H035 | -22.72 | H107 | -6.20 | H019 | 6.17 |
| H121 | -22.48 | H074 | -6.10 | H028 | 6.18 |
| H062 | -21.91 | H053 | -5.33 | H012 | 6.30 |
| H067 | -21.81 | H003 | -5.09 | H093 | 6.52 |
| H091 | -20.87 | H040 | -4.96 | H027 | 6.70 |
| H008 | -20.05 | H097 | -4.65 | H094 | 7.17 |
| H113 | -19.35 | H042 | -3.53 | H075 | 7.68 |
| H124 | -19.00 | H101 | -3.40 | H058 | 8.04 |
| H010 | -18.86 | H085 | -2.84 | H002 | 8.07 |
| H120 | -18.49 | H083 | -2.08 | H049 | 8.80 |
| H054 | -18.36 | H014 | -2.02 | H117 | 8.85 |
| H126 | -16.55 | H068 | -1.82 | H125 | 8.96 |
| H105 | -15.81 | H004 | -1.45 | H023 | 9.38 |
| H022 | -15.46 | H011 | -1.40 | H063 | 9.54 |
| H076 | -15.45 | H036 | -0.91 | H031 | 9.76 |
| H037 | -14.88 | H122 | -0.80 | H009 | 10.09 |
| H087 | -14.83 | H051 | -0.29 | H064 | 10.30 |
| H110 | -14.51 | H079 | 0.18 | H001 | 11.08 |
| H050 | -14.43 | H060 | 0.18 | H015 | 12.76 |
| H016 | -14.33 | H073 | 0.41 | H006 | 14.37 |
| H052 | -13.13 | H089 | 0.45 | H029 | 14.41 |

| | | | | | |
|---|---|---|---|---|---|
| H065 | -12.88 | H115 | 0.70 | H043 | 15.08 |
| H066 | -12.37 | H032 | 0.73 | H116 | 15.34 |
| H005 | -11.78 | H030 | 0.96 | H047 | 16.94 |
| H013 | -11.55 | H045 | 1.66 | H007 | 18.35 |
| H072 | -11.24 | H021 | 1.91 | H080 | 18.51 |
| H055 | -11.05 | H084 | 2.11 | H039 | 19.89 |
| H086 | -9.91 | H103 | 2.15 | H096 | 30.77 |
| H059 | -9.79 | H070 | 2.28 | H102 | 35.46 |
| H077 | -9.46 | H106 | 2.49 | MEAN | -3.14 |
| H071 | -9.14 | H095 | 2.53 | | |

Supplementary Table S13. Summary of the linkage maps of the segregating population 'Candonga' x 'Senga Sengana'. Some linkage groups which could not be linked by markers were split into several parts (1, 2 and sometimes 3, 4).

| Female map | | | | Male map | | | |
|---|---|---|---|---|---|---|---|
| LG | Markers | Size (cM) | Markers/cM | LG | Markers | Size (cM) | Markers/cM |
| 1A | 322 | 48 | 0.15 | 1A | 92 | 27.5 | 0.3 |
| 1B | 135 | 19.9 | 0.15 | 1B | 247 | 54.8 | 0.22 |
| 1C | 215 | 65.3 | 0.3 | 1C | 229 | 59.1 | 0.26 |
| 1D | 144 | 65.1 | 0.45 | 1D | 125 | 52.7 | 0.42 |
| 2A | 389 | 72 | 0.19 | 2A | 61 | 54.2 | 0.89 |
| 2B1 | 66 | 21.2 | 0.32 | 2B | 349 | 79.4 | 0.23 |
| 2B2 | 40 | 4.5 | 0.11 | 2C1 | 146 | 49.2 | 0.34 |
| 2C | 304 | 83.3 | 0.27 | 2C2 | 9 | 50.6 | 5.62 |
| 2D | 317 | 67.1 | 0.21 | 3A | 170 | 76.2 | 0.45 |
| 3A | 419 | 109.7 | 0.26 | 3B | 163 | 74.3 | 0.46 |
| 3B | 114 | 56.7 | 0.5 | 3C | 241 | 71.1 | 0.3 |
| 3C | 270 | 85.3 | 0.32 | 3D | 228 | 58 | 0.25 |
| 3D1 | 207 | 47.2 | 0.23 | 4A1 | 121 | 67.1 | 0.55 |
| 3D2 | 14 | 0.9 | 0.07 | 4A2 | 7 | 9.3 | 1.32 |
| 4A | 159 | 41 | 0.26 | 4B | 147 | 57.7 | 0.39 |
| 4B | 127 | 23.6 | 0.19 | 4C | 137 | 57.7 | 0.42 |
| 4C | 239 | 80.6 | 0.34 | 4D | 187 | 50.1 | 0.27 |
| 4D | 218 | 68.3 | 0.31 | 5A1 | 80 | 21.2 | 0.26 |
| 5A1 | 471 | 72.9 | 0.15 | 5A2 | 17 | 3.6 | 0.21 |
| 5A2 | 210 | 62.9 | 0.3 | 5B | 86 | 51.1 | 0.59 |
| 5C1 | 193 | 14.7 | 0.08 | 5C | 242 | 56.7 | 0.23 |
| 5C2 | 90 | 31 | 0.34 | 5D | 105 | 70.6 | 0.67 |
| 5D1 | 172 | 74.1 | 0.43 | 6A | 216 | 86.4 | 0.4 |
| 5D2 | 26 | 33.5 | 1.29 | 6B | 457 | 82.3 | 0.18 |

| | | | |
|---|---|---|---|
| 5D3 | 23 | 20.3 | 0.88 |
| 6A1 | 182 | 133 | 0.73 |
| 6A2 | 230 | 123.4 | 0.54 |
| 6B1 | 31 | 43.7 | 1.41 |
| 6B2 | 69 | 13.6 | 0.2 |
| 6B3 | 43 | 19.1 | 0.44 |
| 6C1 | 33 | 106.5 | 3.23 |
| 6C2 | 11 | 106.5 | 9.68 |
| 6C3 | 62 | 106.5 | 1.72 |
| 6D1 | 102 | 27.4 | 0.27 |
| 6D2 | 39 | 36.8 | 0.94 |
| 7A1 | 105 | 100.4 | 0.96 |
| 7A2 | 145 | 54.6 | 0.38 |
| 7B | 469 | 42.5 | 0.09 |
| 7C | 155 | 33.2 | 0.21 |
| 7D1 | 63 | 39.1 | 0.62 |
| 7D2 | 49 | 14.9 | 0.31 |
| 7D3 | 40 | 21.5 | 0.54 |
| 7D4 | 65 | 6.3 | 0.1 |
| Total | 6777 | 2298.5 | |

| | | | |
|---|---|---|---|
| 6C | 432 | 78 | 0.18 |
| 6D | 321 | 44.8 | 0.14 |
| 7A | 150 | 39.1 | 0.26 |
| 7B | 148 | 13.9 | 0.09 |
| 7C1 | 271 | 65.1 | 0.24 |
| 7C2 | 50 | 24.9 | 0.5 |
| 7C3 | 8 | 10.1 | 1.26 |
| 7D | 175 | 56.5 | 0.32 |
| Total | 5417 | 1653.1 | |

Figure S1

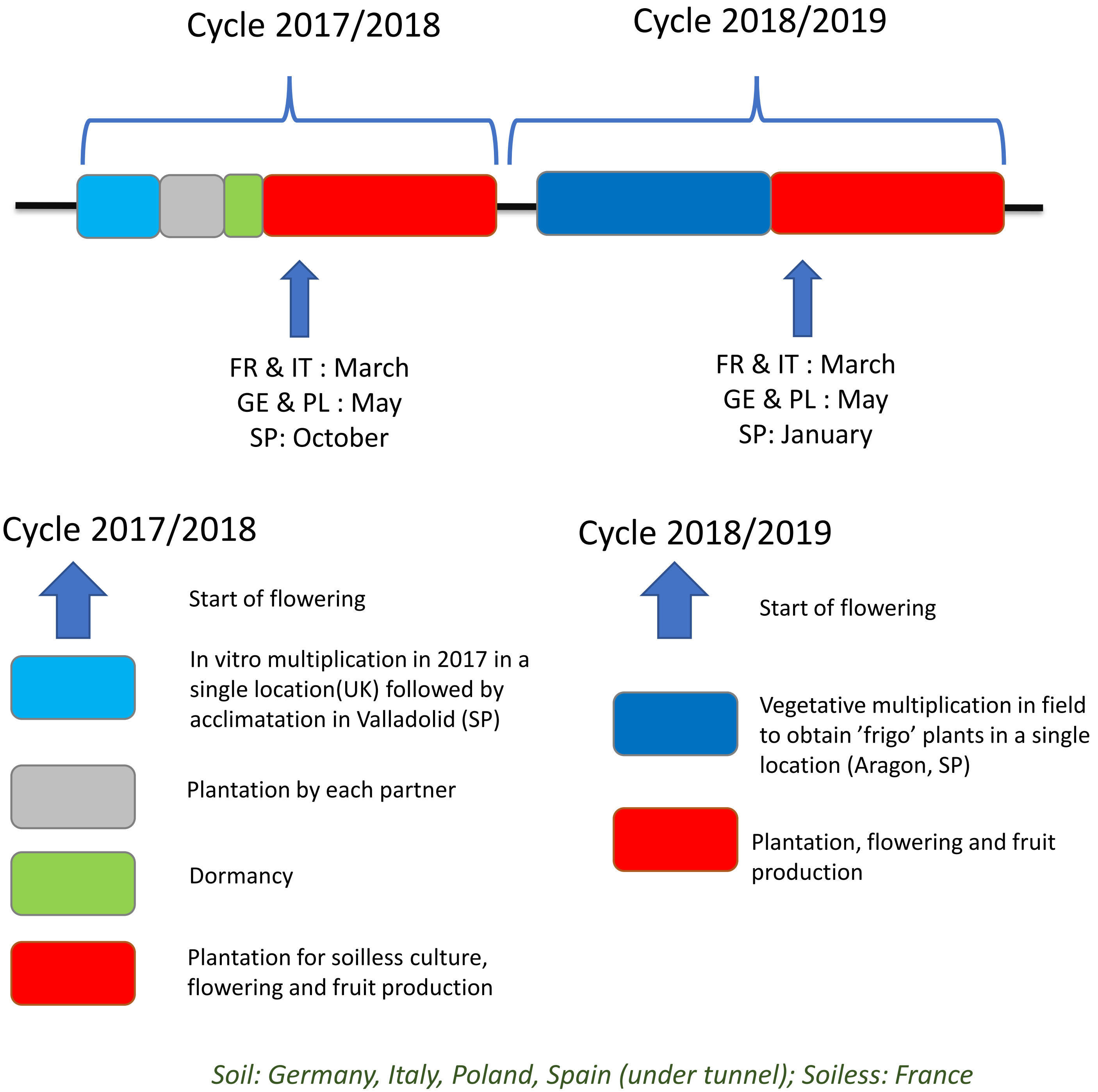

Supplementary Figure S1. Culture workflow of the European Goodberry project. Plants were raised in a single location (Valladolid, Spain in 2017 and Aragon in 2018). In 2017 vitro plants were sent to each partner for acclimatization before planting. In 2018, 'frigo' plants (cold stored plants) were sent to each partner for planting. SP, Spain; IT, Italy; FR, France; GE, Germany; PL, Poland.

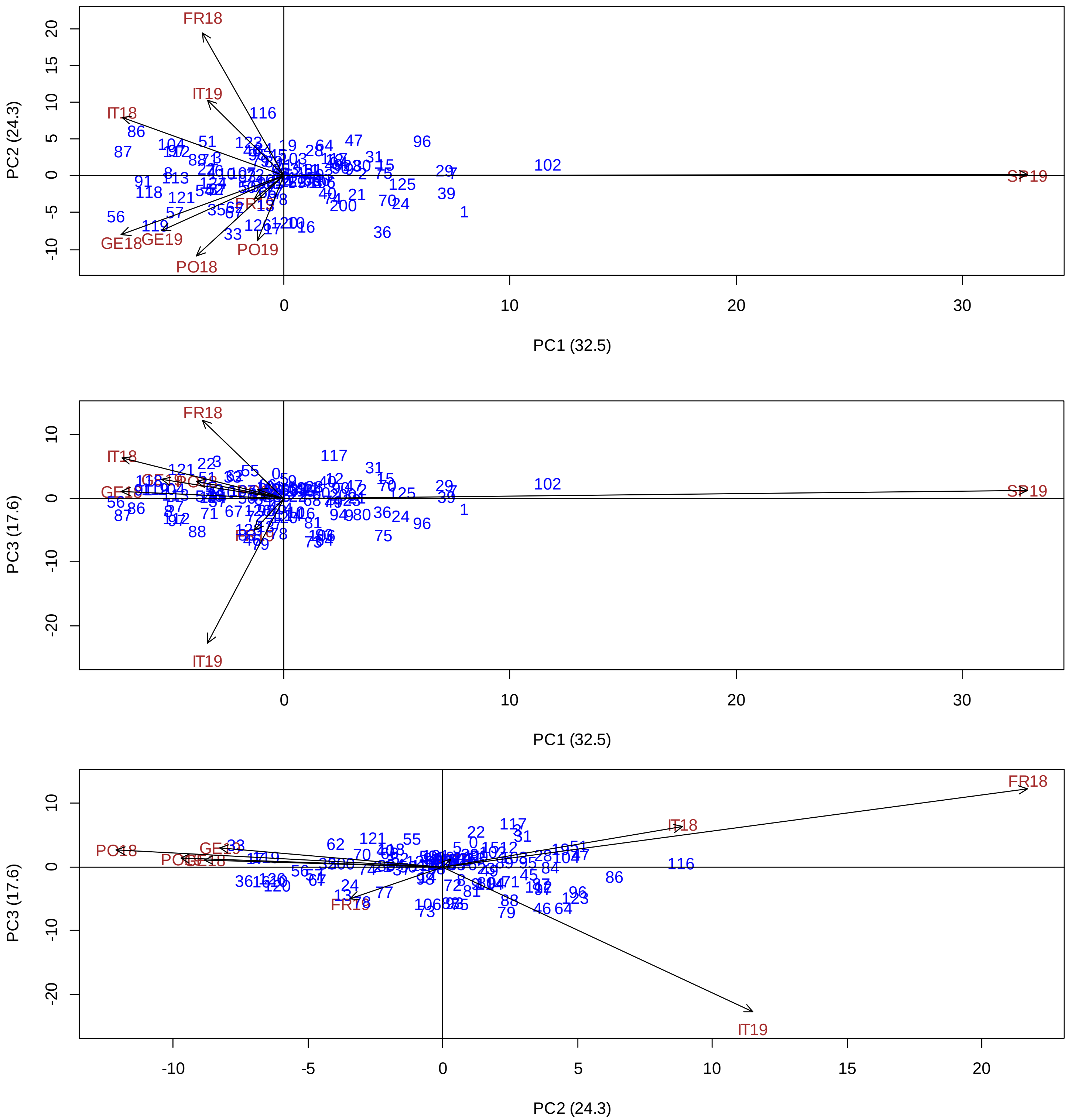

Supplementary Figure S2. Biplots of the first three components of AMMI analysis of flowering time. Analysis was performed for the nine environments and flowering time was expressed as GDD. The blue numbers correspond to individuals of the ‘Candonga’ x ‘Senga Sengana’ population. Environments are represented by the dark lines and are indicated by the red codes.

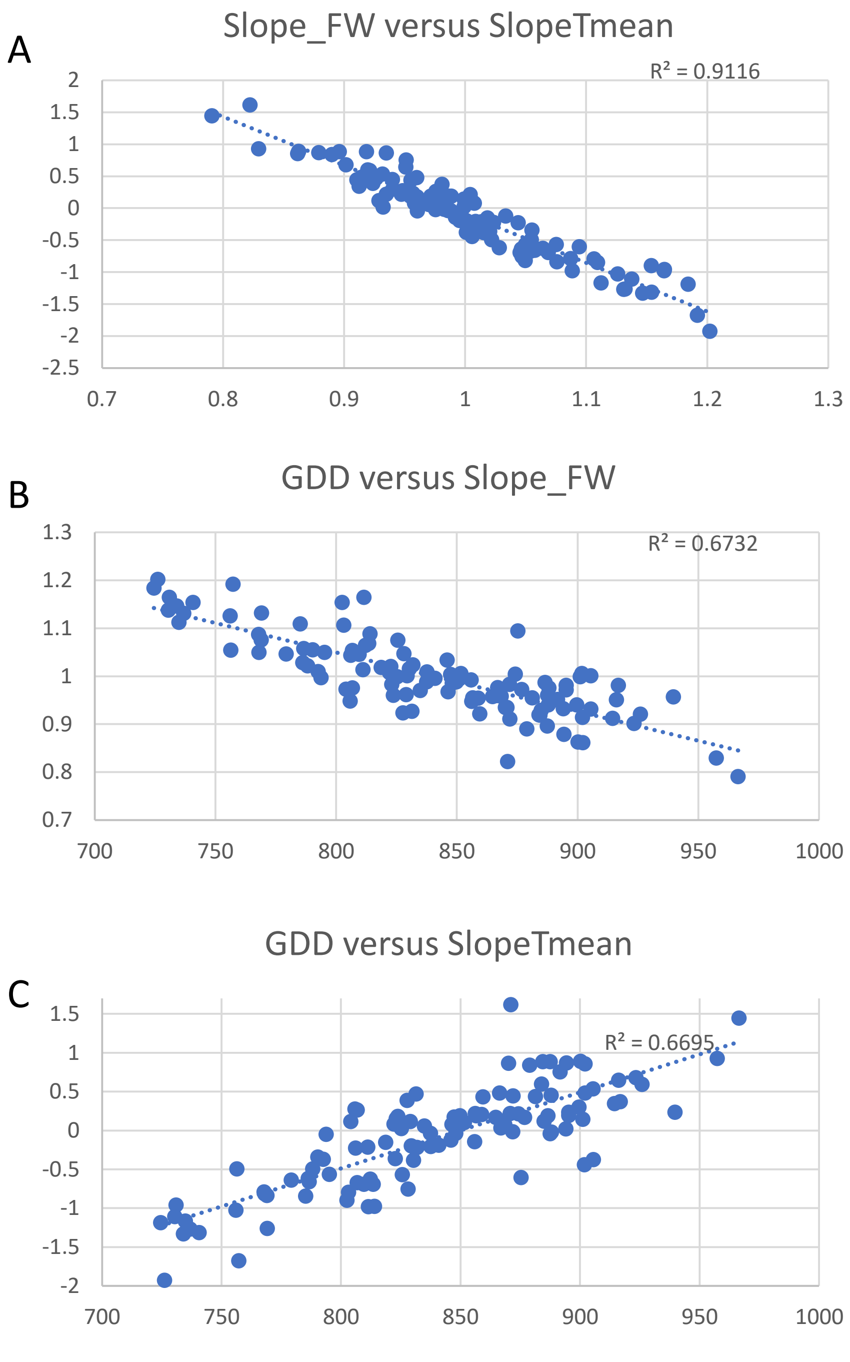

Supplementary Figure S3. Relationship between GDD, slope_FW and slope_Tmean. Slope_FW value represents the linear regression of each genotype on the nine environmental means as proposed by Finlay-Wilkinson (FW). SlopeTmean represents the linear regression of each genotype on the nine environments characterized by their Tmean as propose by factorial regression.

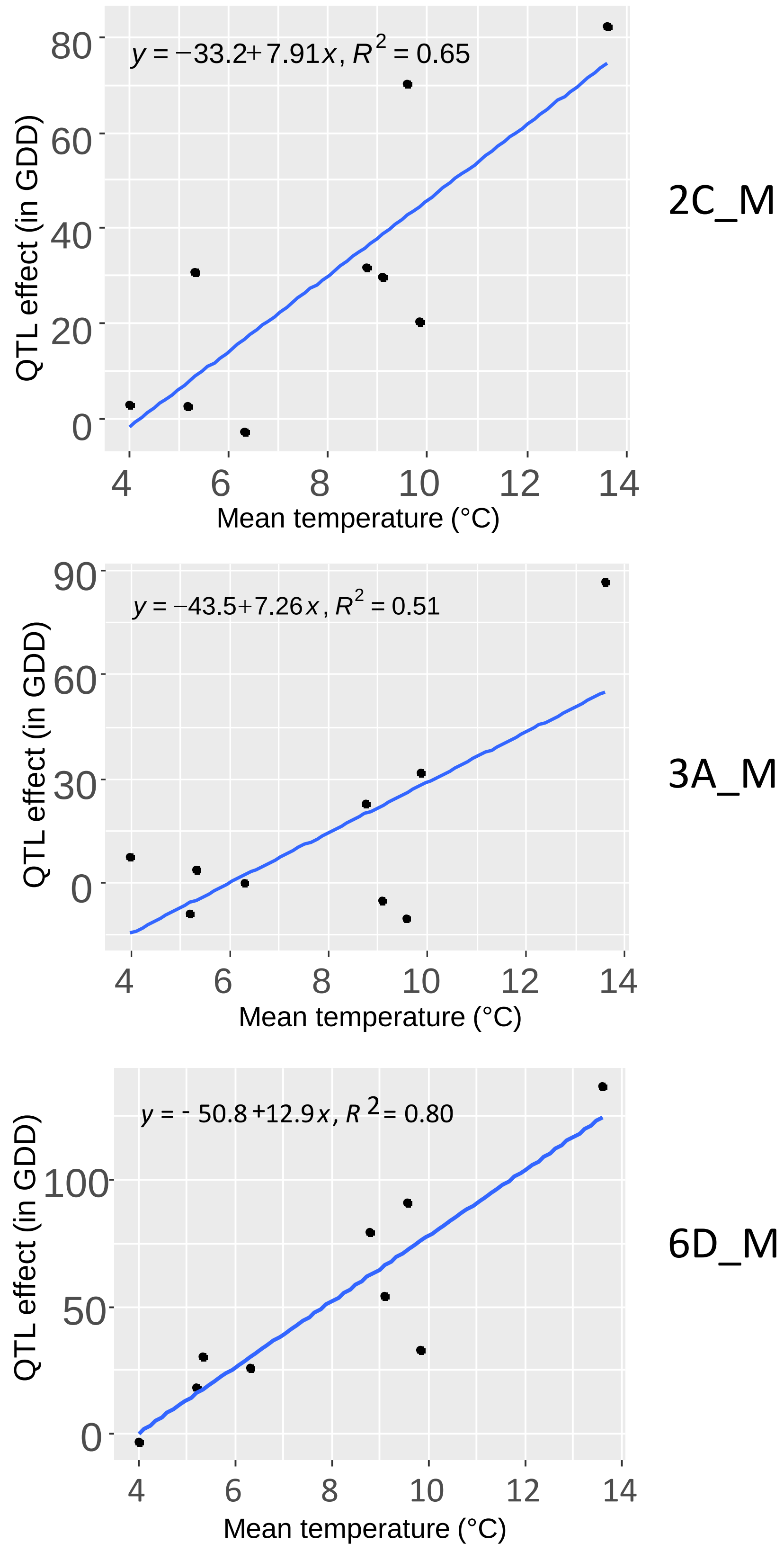

Supplementary Figure S4. Relationship between the QTL effect expressed in GDD and the temperature. Mean temperature was calculated from the 1st of January to the end of flowering in each location.

Supplementary Figure S5. Single marker analysis showing the effect of the Affymetrix® allele linked to the flowering time QTL.

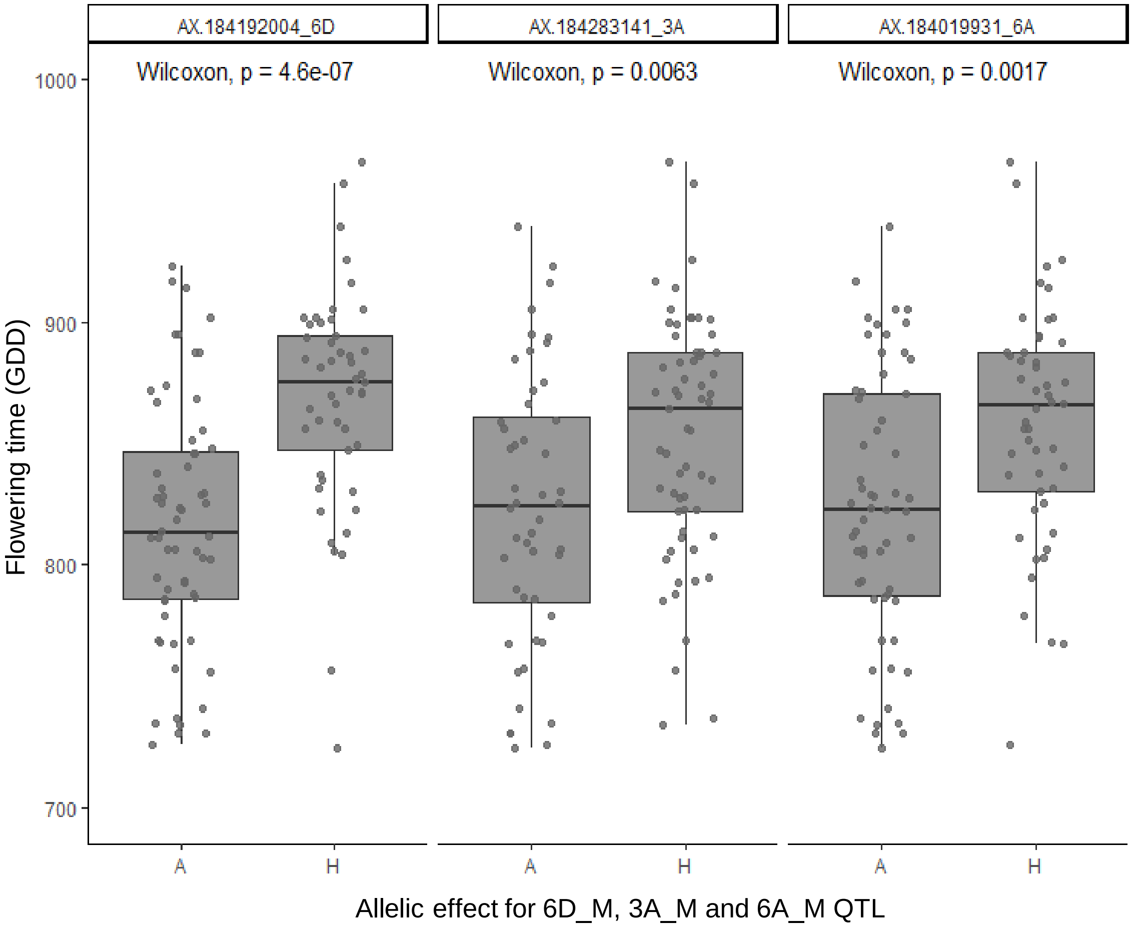

Supplementary Figure S5. Single marker analysis showing the effect of the Affymetrix® allele linked to the flowering time QTL. Wilcoxon test.
